# The neurocomputational mechanisms of hierarchical linguistic predictions during narrative comprehension

**DOI:** 10.1101/2025.03.27.645665

**Authors:** Faxin Zhou, Siyuan Zhou, Yuhang Long, Adeen Flinker, Chunming Lu

**Affiliations:** State Key Laboratory of Cognitive Neuroscience and Learning and IDG/McGovern Institute for Brain Research, Faculty of Psychology, Beijing Normal University, No. 19 Xinjiekouwai Street, Beijing 100875, PR China; Biomedical Engineering Department, Tandon School of Engineering, New York University, Brooklyn, 11201, NY, USA; Institute of Developmental Psychology, Faculty of Psychology, Beijing Normal University, Beijing, 100875, China; Neurology Department, Grossman School of Medicine, New York University, New York, 10016, NY, USA

**Keywords:** Narrative comprehension, Prediction hierarchy, Information updating, fMRI, Neural computation

## Abstract

Language comprehension requires a listener to predict the upcoming inputs of linguistic units with multiple timescales based on previous contexts, but how the prediction process is hierarchically represented and implemented in the human brain remains unclear. Combining the natural language processing (NLP) approach and functional Magnetic Resonance Imaging (fMRI) in a narrative comprehension task, we first applied the group-based general linear model (gGLM) to identify the neural underpinnings associated with the language prediction on word and sentence. Our results revealed a cortical architecture supporting the prediction, extending from the superior temporal cortices to the regions in the default mode network. Then, we investigated how these adjacent levels interact with each other by testing two rival hypotheses: the continuous updating hypothesis posits that the higher level of the representational hierarchy is continuously updated as inputs unfold over time, while the sparse updating hypothesis states that the higher level is only updated at the end of their preferred timescales of linguistic units. By conducting computational modeling and autocorrelation analysis, we found the sparse model outperformed the continuous model and the updating might occur at the sentence boundaries. Together, our results extend the linguistic prediction from the small timescales such as words to large timescales such as sentences, providing novel insights into the neurocomputational mechanisms of information updating within the linguistic prediction hierarchy.

## Introduction

Language comprehension requires a listener to predict the upcoming input based on previous knowledge and contexts [1–4]. Prediction can reduce the computational load in the brain [5], enabling the listener to instantaneously process the highly dynamic speech flow (2-5 words per second) in the natural language [6,7]. Despite converging studies suggesting that natural language is hierarchically organized with temporally smaller linguistic units embedded in the larger linguistic units [8–11], it remains unclear how our brain represents and implements the linguistic prediction of multilevel units with different timescales within such a hierarchical structure.

Previous studies on language prediction have primarily concentrated on the neural representations related to the upcoming linguistic units at smaller timescales, such as phonemes and words. Evidence along this line of research has shown that phoneme prediction is represented in the bilateral primary auditory cortices [1,12], while word prediction is represented in a relatively distributed brain network including the bilateral STG, left inferior parietal lobule (IPL), bilateral inferior frontal gyrus (IFG) and bilateral dorsolateral PFC (dlPFC) [13,14]. However, it remains to be determined how the linguistic prediction is conducted in the human brain across multiple timescales, particularly for those with longer timescales. As a larger linguistic unit than word and phoneme, sentence can convey more complex concepts and meanings [15,16]. The rich context embedded in sentences also enables individuals to navigate complex communication situations [17,18], such as attuning to social-emotional cues [19] and inferring underlying communicative intentions [20]. Moreover, the investigation of sentence prediction can provide a detailed understanding on narrative language comprehension for both human brain and artificial intelligence (AI) systems [21,22].

Therefore, the comparison between the predictive representations of larger-timescale units such as sentences and smaller-timescale units such as words are vital for understanding the unique contributions of multiscale linguistic units within the prediction hierarchy.

Additionally, it is also crucial to understand how information updates between adjacent levels within the prediction hierarchy. As the neural representation of the prediction hierarchy has not been completely depicted yet, only a few studies have tested this issue and primarily focused on the lower levels [1,8,12,23]. A debate emerged from these studies concentrates on how information is updated along the hierarchy. One perspective suggests that higher levels are updated continuously as inputs from lower levels unfold over time (i.e., continuous updating hypothesis). For example, studies on auditory perception [24] and narrative comprehension [25] have shown that neural responses at higher levels have a gradual rather than an abrupt pattern of increase when new inputs appear. Moreover, computational models built on the continuous updating hypothesis can well capture the neural features during context constructing and forgetting [25]. A rival perspective suggests that higher levels are only updated at the end of their preferred timescales, thus showing an abrupt rather than a gradual increase regarding the neural responses (i.e., sparse updating hypothesis). For example, evidence shows that the neural responses of the precuneus changed sharply at its preferred timescales (i.e. the event boundaries) [26]. Additionally, a study using the recurrent neural network (RNN) model showed that the sparse updating model could differentiate the processing architecture of the human brain along the temporo-parietal axis, while the continuous updating model could not [23]. Based on these perspectives, in this study, we would test which model (the continuous or the sparse model) could best explain the mode of information update from the lower-level brain regions representing the word prediction to the higher-level brain regions representing the sentence prediction.

To investigate the prediction hierarchy and test the information-updating modes in the human brain, we combined the natural language processing (NLP) approach and the neural computational modeling method to analyze brain signals from individuals engaged in a narrative comprehension task, recorded using functional Magnetic Resonance Imaging (fMRI). In this task, 31 participants were asked to listen to three aurally presented stories that were played either forward or backward. The forward condition intrinsically contains a hierarchy for the linguistic prediction [8,9], while the backward condition was used as a control for acoustic features. Next, we aimed to quantify the predictive relationship between the previous contexts and the upcoming linguistic units at both word and sentence levels. While decoder-only transformer architectures (e.g., generative pre-trained transformer (GPT) model) have been widely used in prior studies as they are conceptually aligned with the idea of linguistic prediction [1,13], their autoregressive property primarily manifests at smaller timescales (e.g., words), and it remains challenging to utilize them for investigating predictive mechanism of large units such as sentences. Therefore, to overcome this limitation, we employed a multiple ridge regression modeling procedure to capture the predictive relationship at multiple timescales (Fig. 1a), by leveraging the linguistic embeddings obtained from Bidirectional Encoder Representations from Transformers (BERT) model [27,28]. Note that the BERT model employed in this step was only for generating linguistic representations rather than simulating the linguistic prediction process. After obtaining the predictive representations, we further identified the neural underpinnings of multilevel linguistic prediction using the group-based general linear model (gGLM). Finally, to distinguish continuous/sparse updating hypotheses and explore the temporal dynamics across word and sentence levels, we built a set of neural computational models and tested them by simulating the fMRI responses.

**Figure 1.**
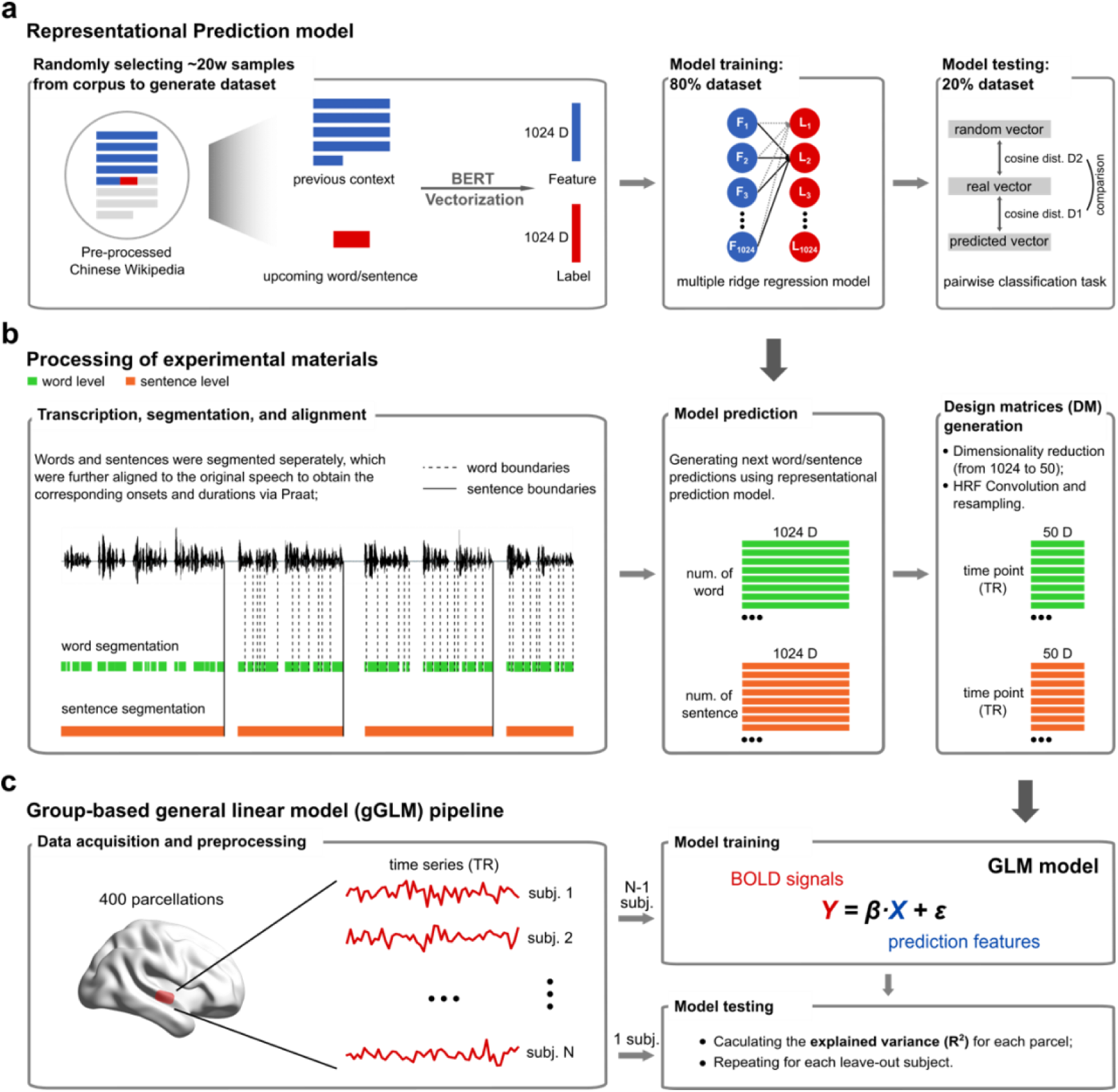
Schematic demonstration of the analytic approach. a) Training and testing the multiple ridge regression models. The dataset (∼0.2 million samples) was generated from Chinese Wikipedia, by randomly selecting the prior linguistic context and upcoming linguistic units (word or sentence). The context and linguistic unit were transformed into fixed-length vectors via the WWM-RoBERTa model. Then, 80% of samples were used to train a multiple ridge regression model to capture the predictive relationship between the context and linguistic unit, and the rest 20% was used for model evaluation. Word and sentence prediction models were trained separately. b) Processing of the experiment materials. The story audios were transcribed, segmented, and aligned at both word (via “jieba” toolbox implemented in Python) and sentence level (via sentence boundary segmentation task), which were further used to generate the predictive representations using the regression models. These representations were further reduced to 50 dimensions, resampled, and convolved with the hemodynamic response function (HRF) for further analyses. c) Roadmap of the group-based general linear model (gGLM). BOLD signals were collected while participants listened to stories. The BOLD signals were pre- processed and parceled to 400 according to Schaefer et al. (2018). On each parcel, a leave-one-subject-out (LOSO) cross-validation approach was employed to obtain the explained variance (*R^2^*) for each parcel across participant.

In line with the prior evidence, we first hypothesized that the word prediction would be primarily located in lower-order brain regions such as STG [13,14]. Additionally, given recent advances showing that default mode network (DMN) is largely employed in the anticipation of future events (also known as prospective memory) [29–32] and actively involved in narrative understanding (especially at longer timescales) [10,11,32–34], we expected to observe neural representations of sentence prediction within DMN regions. Furthermore, considering the chunking property of the language system [8,9], we postulated that the functional interactions between these two levels would be conducted in a sparse manner, which has been shown to be more computationally efficient and capable of accelerating processing speed [23,35]. Overall, we provided novel evidence for the prediction hierarchy in the brain and the information updating strategy regarding cross-level interactions.

## Results

### Behavioral performance in narrative comprehension

In the narrative comprehension task, participants were requested to passively listen to the three stories while fixating on a cross at the center of the black screen. The sequence of the six audios (three in the forward and three in the backward conditions) was counterbalanced across participants. An interval with flexible length was inserted between two audios. The details of the story stimuli can be found in Table 1.

**Table 1.** Descriptive statistics of the three stories.

|  | Number.<br>of<br>characters | Total<br>duration<br>(min:sec) | Words |  |  | Sentences |  |  |
| --- | --- | --- | --- | --- | --- | --- | --- | --- |
|  |  |  | No. | Mean<br>duration | *S.D.* | No. | Mean<br>duration | *S.D.* |
| Story 1 | 959 | 3:00 | 533 | 0.30 | 0.14 | 30 | 5.62 | 3.22 |
| Story 2 | 1387 | 4:45 | 814 | 0.28 | 0.15 | 40 | 6.48 | 2.13 |
| Story 3 | 1475 | 6:18 | 831 | 0.34 | 0.18 | 53 | 6.19 | 3.07 |

Note: *S.D.* represents standard deviation.

At the end of each story, the participants were tested on how well they perceived and comprehended this story (see Materials and Methods). To test whether the participants perceived the audio stimuli clearly, one sample *t*-test was conducted on the perception scores against the chance level. The results showed that all the perception scores were significantly higher than the chance level (*ps* < 0.05, Table 2). Additionally, an ANOVA was performed on the scores across three stories to test their comparability. The results did not show any significant differences among three stories regarding clarity (*F*(2, 90) = 0.203, *p* = 0.817, *f* = 0.081), familiarity (*F*(2, 90) = 0.594, *p* = 0.554, *f* = 0.138), and complexity (*F*(2, 90) = 3.000, *p* = 0.055, *f* = 0.311). These results ensured the reliability of the comprehension scores reported below.

**Table 2.** Participants’ performances for all three stories.

|  | Perception score |  | Comprehension score |  |
| --- | --- | --- | --- | --- |
| | Mean | $S.D.$ | Mean | $S.D.$ |
| Story 1 | 4.226* | 0.805 | 4.709* | 0.783 |
| Story 2 | 4.323* | 0.599 | 3.000* | 0.000 |
| Story 3 | 4.323* | 0.653 | 2.839* | 0.374 |

Note: $S.D.$ , standard deviation. \*, $p < 0.05$ .

Next, we tested how well the participants comprehended the stories. A Wilcoxon signed rank test was conducted on scores against the chance level and a Kruskal-Wallis test was performed on scores across three stories. Non-parametric tests were selected here because of the non-normality of the data (see Materials and Methods). The results showed that the comprehension score of each story was significantly higher than the chance level (*ps* < 0.05, Table 2) but did not significantly differ among three stories (Kruskal-Wallis test, *H*(2) = 5.524, *p* = 0.063). Thus, the comprehension scores were summed across the three stories to index the overall performance (mean across participants = 10.548, *S.D.* = 0.850), which was also significantly higher than the chance level (i.e., 5.5) (Wilcoxon signed rank test, *T*(31) = 496.000, *p* < 0.001). It should be noted that although there were marginally significances for story complexity and comprehension scores, we actually analyzed these stories separately in the subsequent analyses and trained the encoding model with the leave-one-subject-out (LOSO) method. Therefore, the difference between stories would not impact the results.

### The predictive representations of words and sentences

To determine the cortical architecture supporting the prediction hierarchy, we needed to obtain the predictive representation of sentences and replicate previous findings on the predictive representation of words. To this end, we employed a large language model (LLM) approach that ensured the comparability of results of word and sentence prediction.

First, the Robustly Optimized Bidirectional Encoder Representations from Transformers (BERT) Pretraining Approach with Whole Word Masking (WWM-RoBERTa) model was employed to vectorize the language inputs [36]. WWM-RoBERTa is a variant of the BERT model, with a bigger architecture, larger batch size, and training dataset [37]. More importantly, it is trained on the prediction task of whole words rather than characters, hence possessing higher generalizability and adaptability for Mandarin [38]. Previous studies in both NLP and cognitive neuroscience have shown that BERT models are very cognizant of grammatical and discourse markers [27,28] and can provide better representations of language information than other models [37,39–41]. Practically, word representations were obtained by inputting each word solely into the BERT model (without contexts), and sentence/context representations were acquired by taking the average of all word embeddings within the text (see Methods).

Second, the multiple ridge regression modeling method was used to capture the predictive relationship between the prior linguistic contexts and the prediction targets (i.e., the upcoming linguistic units) based on the vectorial representations. The ridge regression model is applied by assuming that the prediction relationship becomes linear after feature extraction from the BERT model. This assumption is based on many of previous studies showing that, after the feature engineering of the large language models (LLMs), the language embeddings show the analogical relations (e.g., *queen – woman ≈ king - man*) and exhibit seemingly linear behavior [42–44]. Moreover, the ridge regression model can also effectively mitigate the overfitting problem. In practice, each dimension of the target vectors was predicted using a different ridge regression model, with parameters estimated from training data (80%) and validated from the test data (20%) (Fig. 1a, middle panel; see Materials and Methods). Regression models were constructed for word and sentence prediction respectively.

A pairwise classification task was employed to evaluate the performance of two ridge regression models (Fig. 1a, right panel) [45]. In this task, we calculated cosine distances between the vectors predicted by the models and the real vectors of the targets (denoted as D1). D1 was compared with the cosine distances between the predicted vectors and the vectors that were randomly selected from the test set (denoted as D2). We took D2 as the baseline because the random vectors represent a scenario without any predictive relationship, given that the distributions of D2 is centered around 1 (Fig. 2a-b, gray histograms). We first randomly selected 1000 samples from the testing set and found the cosine distances between the real and predicted targets were significantly lower than that between the random and predicted targets at both word and sentence levels (paired two-sample *t*-test; word level: *t*(999) = 19.18, *p* < 0.001, *d* = 0.876; sentence level: *t*(999) = 43.870, *p* < 0.001, *d* = 1.439; Fig. 2a, b). Additionally, we calculated the Pearson correlation between predicted and real targets (*r1*) or random-generated targets (*r2*) to supplement the above results. The results also showed significant differences between these two conditions at both word and sentence levels (paired two-sample *t*- test after Fisher-z transformation on the *r* values; word level: *r1* = 0.078 ± 0.112; *r2* = -0.001 ± 0.065; *t*(999) = -18.264, *p* < 0.001, *d* = 0.876; sentence level: *r1* = 0.113 ± 0.112; *r2* = 0.004 ± 0.065; *t*(999) = -43.812, *p* < 0.001, *d* = 1.439; Fig. S1a).

**Figure 2.**
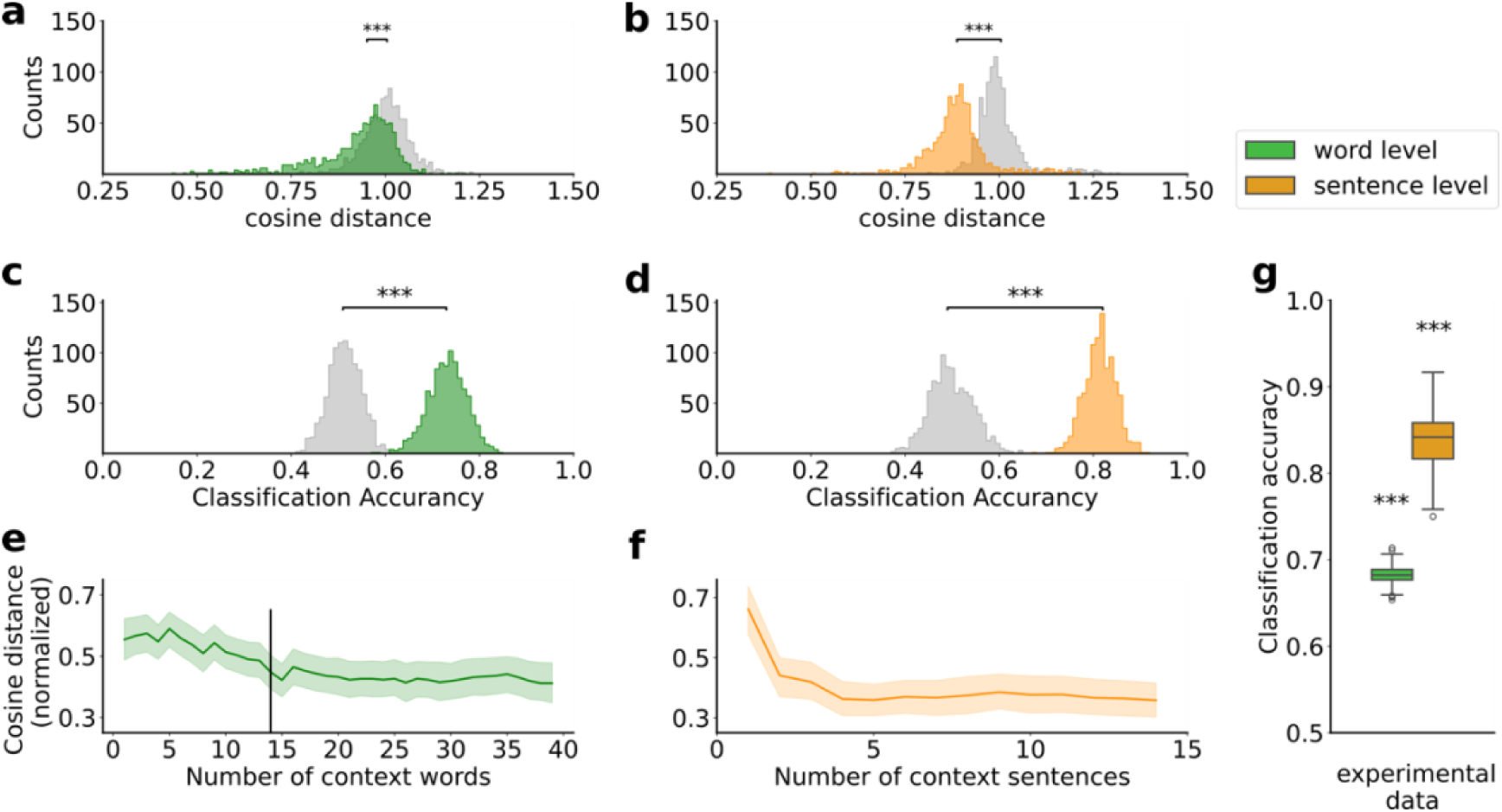
Results of the representational prediction models’ performances. a, b) The cosine distances between predicted targets and the real ones for word (green) and sentence (orange) prediction models against the cosine distances between predicted targets and the random vectors (gray). c, d) The classification accuracies of both word (73.234 ± 4.265%, green) and sentence predictions (81.471 ± 3.478%, orange) are higher than the random data (gray). e, f) Both models showed the incremental context effect. The cosine distances were max-min normalized, and the vertical line in e) represents the mean number of words in a sentence. g) The model performances of experimental materials.

Third, we replicated the above procedure 1000 times for robustness. If D1 was smaller than D2, the classification result was coded as correct; otherwise, the result was coded as incorrect. The results showed significantly higher prediction accuracy than the chance level (i.e., 50%) for both word (73.234 ± 4.265%, *p* < 0.001; Fig. 2c) and sentence (81.471 ± 3.478%, *p* < 0.001; Fig. 2d) models. To validate this result, we randomly shuffled the pairwise relationship between the prediction targets and the prior linguistic contexts, whereby a random dataset was generated. Then, the pairwise classification task was applied to this random dataset. The results showed that the classification accuracy in the random dataset did not significantly differ from the chance level (1000 times permutation test; word model: 50.132 ± 3.458%, *p* = 0.351; sentence model: mean = 49.812 ± 4.719%, *p* = 0.503; Fig. 2c, d). Moreover, the classification accuracy in the original dataset was significantly higher than that of the random dataset (two-sample *t*-test; word model: *t*(1998) = 128.800, *p* < 0.001, *d* =5.760; sentence model: *t*(1998) = 170.788, *p* < 0.001, *d* = 7.638; Fig. 2c, d).

Finally, two ridge regression models were further tested on the narratives used in the present study. The results showed that the classification accuracy was 68.241% (*S.D.* = 0.873%) for the word model and 83.731% (*S.D.* = 2.811%) for the sentence model, respectively, both of which were significantly higher than the chance level (1000 times permutation test; word level: *p* < 0.001; sentence level: *p* < 0.001; Fig. 2g).

Together, these results indicated that the above models could robustly replicate the predictive relationship between the prior linguistic contexts and the target words. Moreover, we additionally provided novel results for the sentence prediction. We observed that the sentence model outperformed the word model on both the corpus dataset and our experimental materials. We reasoned that this result may stem from the effective BERT representation of sentence, which contains more dynamic, context-dependent information. Furthermore, when computing sentence embeddings, we averaged word embeddings across words of a sentence, which might also increase the signal-to-noise ratio (SNR) compared to the computation at the word level. However, we suggested that absolute model accuracy is not a suitable indicator of predictive quality, as our models are designed to predict the latent space of BERT features. Instead, the overall statistical significance of model performance better reflects the model’s capability in capturing predictive relationships between the contexts and the targets within the language corpus.

### Validation of the regression models based on the incremental context effect

Previous evidence has indicated that the prediction of the upcoming linguistic unit is incrementally influenced by the prior linguistic context. Thus, to be biologically plausible, the aforementioned models should “behave” as a function of the length of the prior context, i.e., the incremental context effect [13,41]. We tested this account by gradually increasing the inputs (i.e., number of words/sentences) of the prior linguistic context. The results showed that, for the word model, the cosine distance between the predicted vectors and the real vectors decreased as the number of words prior to the prediction target increased.

Next, the knee point was identified through kneed python toolbox (https://github.com/arvkevi/kneed), which applies a rotation-based algorithm to find the points with the maximum curvature [47]. The knee point of the word model appeared at the boundary of a sentence (Fig. 2e). We determined the mean lengths of a sentence by averaging the length of all sentences in Chinese Wikipedia (approximately 3.9 million sentences, median length is 15 words, Fig. S1b).

Similarly, for the sentence model, the cosine distance between the predicted vectors and the real vectors decreased with the increase in the number of sentences before the prediction targets. A turning point appeared when the number of prior sentences reached 4 (Fig. 2f). These findings confirmed the incremental context effect and further validated our regression models.

### The neural underpinnings of multilevel prediction

We employed the fMRI encoding models to identify the neural underpinnings of the prediction on different linguistic sizes. In previous studies, this method is widely used for its reliability and validity in yielding robust results [41,48–50]. Specifically, in our study, a group-based general linear model (gGLM) was used to relate the pre-whitened BOLD signals to the predicted vectors of words or sentences obtained by two regression models (Fig. 1c, see Materials and Methods). The leave-one-subject-out (LOSO) cross validation approach was employed to avoid overfitting and reduce the non-independence error in the secondary test [51]. Additionally, to improve the computational efficiency, a template with 400 cortical parcels was used for the gGLM analysis [46]. Moreover, to test the concept of “pre-activation” [3] in linguistic prediction, the predicted vectors at time *t* were related to the BOLD signals at time *t*-1. A series of potential confounding factors, such as temporal delays of word/sentence, frequencies of word/sentence usage, and the prior linguistic contexts, were excluded (see Materials and Methods). This analytic procedure was conducted on word and sentence levels separately. Paired two-sample *t*-test was performed between forward and backward conditions on the explained variance (*R^2^*) of the gGLM models. The results were corrected for multiple comparisons using the false discovery rate (FDR) method with a threshold of *p* < 0.01 [52].

At the word level, the results showed that the predictive representations of words were associated with significant activations in the bilateral STG and the upper part of the middle temporal gyrus (MTG) (Fig. 3a, c; Fig. S3a; Table S1). To validate this result, a permutation test was conducted, where the rows and columns of the design matrices in the gGLM analysis were randomly shuffled to remove the predictive relationship between the prior linguistic contexts and the prediction targets. Then, the same gGLM analysis was applied to these shuffled design matrices. This procedure was repeated 1000 times, generating a null distribution of brain activation. The results showed that the real value was significantly higher than the null distribution (*p* < 0.01, FDR corrected; Fig. 3b upper panel, S5a), which confirmed the association between the predictive representations of words and the bilateral STG and MTG.

**Figure 3.**
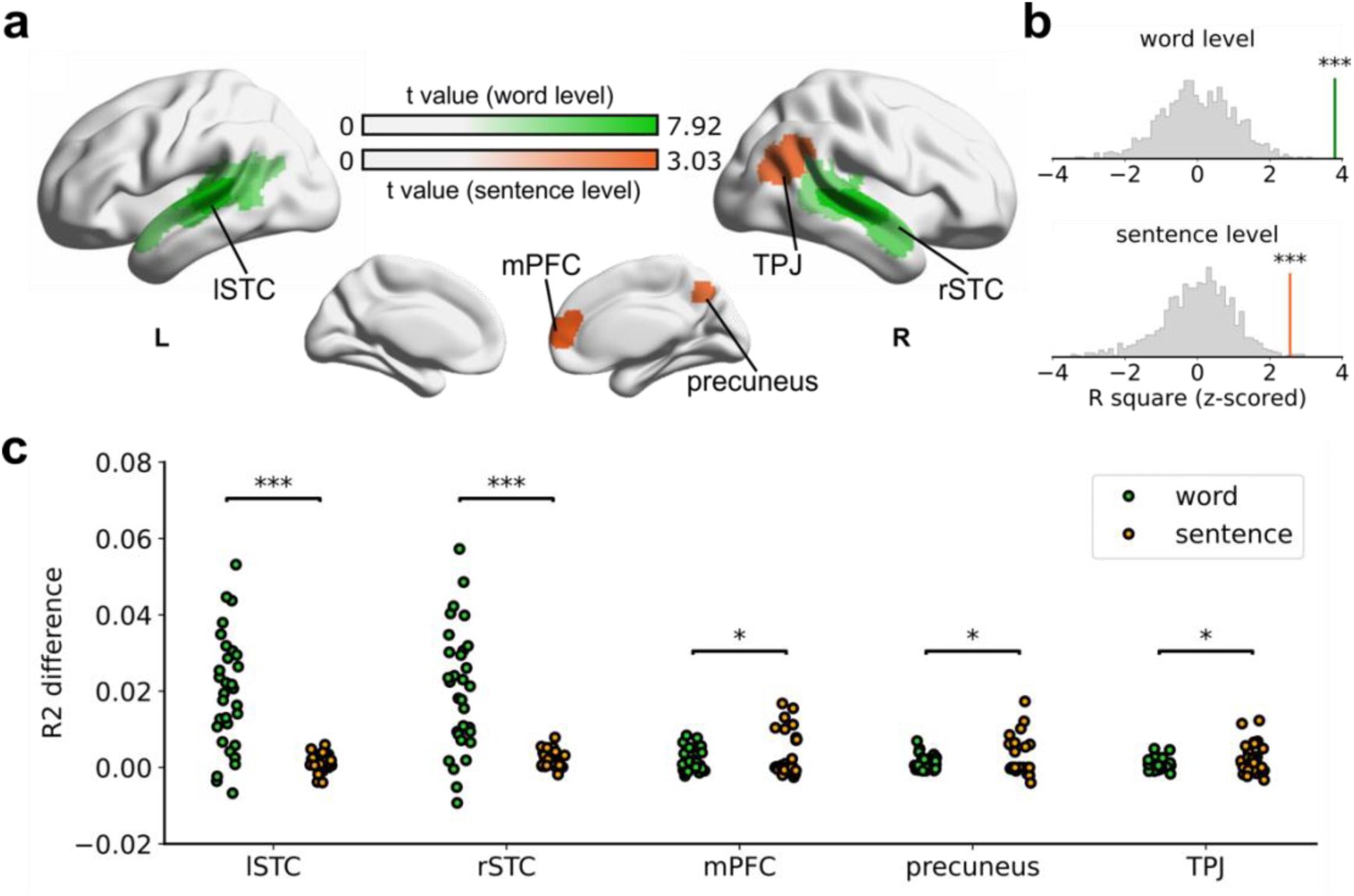
Brain responses associated with predictive representations. a) Brain regions sensitive to predictive representations on different timescales. The results showed the *R^2^* difference between forward and backward conditions, and only significant results are plotted (paired *t*-test, *p* < 0.01, FDR corrected). b) Results of the permutation test. Of all the panels, the x-axis represents the *R^2^* of each permutation, which has been z-scored for display purposes. The gray histograms are the null distributions. The vertical lines indicate the positions of the real values, green for word level and orange for sentence level, respectively. c) the R^2^ difference result for each ROI, where each dot represents one subject. Significant levels are set as *p* < 0.001 (***), *p* < 0.01 (**), *p* < 0.05 (*), and *p* ≥ 0.05 (n.s.).

At the sentence level, the results showed significant activation in the right TPJ, medial PFC (mPFC), and precuneus (Fig. 3a, c; Fig. S3b; Table S1). The same permutation test as aforementioned was performed, which confirmed the significant activation in these brain regions (TPJ: *p* = 0.030; mPFC: *p* = 0.010; precuneus: *p* = 0.010; overall: *p* = 0.010, Fig. 3b lower panel; Fig. S4a; FDR corrected).

We additionally implemented the blockwise permutation test to validate the above results. Following the description of Nichols & Holmes, (2002) and Zhou et al., (2014), we split the BOLD time courses into blocks with each block lasting 10 s, which corresponded to the knee point of our autocorrelation results [53,54] (see below and Fig. 5). We shuffled the blocks in the permutation test and obtained similar results as above (*ps* < 0.01; Fig. S4b).

Furthermore, to differentiate the prediction effect and context effect, we first applied the gGLM modeling pipeline to investigate the context representations at both word and sentence levels. We found that the representations of the past contexts were associated with a widespread distribution of brain regions including the frontal, temporal, and parietal cortices bilaterally (Fig. S5a, b), which was consistent with previous findings on the neural representations of the past linguistic context [10,11,55]. In addition, to detect the unique neural underpinnings of the context effect, we further examined the pure context effects at both word and sentence levels by removing the prediction effect with the variance partitioning (VP) procedures. We found significant representations at both word and sentence levels in the bilateral STG, but the prefrontal cortex was more closely associated with the representation of words (Fig. S5c, d). Please note that the predictive and context features fed into the encoding models are not linearly related due to the non-linear operations used for feature extraction (the Isomap method, see Methods).

Together, these findings suggested that the brain predicted the upcoming words and sentences in a hierarchical manner during language comprehension, and this neural pattern for prediction hierarchy differed from that of context representations.

### Testing the information updating mode across levels of the prediction hierarchy

Next, we aimed to investigate how information updates across the two levels of the prediction hierarchy. To test the sparse and continuous updating models, a series of computational modeling procedures were conducted. This analysis was based on the Predictive Coding (PC) framework [56,57] and used to describe the dynamic interactions between neural underpinnings sensitive to the predictive representations of words and sentences. Briefly, the PC framework posits that the brain employs a deep multi-level cascade when processing the input data, where top-down prediction signals and bottom-up error signals are generated to iteratively update the internal model [58–61]. The theoretical foundation of this framework has empirical support from both computational [57,62,63] and neuroscience studies [56,64].

Specifically, we applied a two-level PC hierarchy to simulate the information update modes. The predictions of words and sentences correspond to the higher and lower levels respectively. According to the PC perspective, the higher level would generate a top-down prediction (*Z*_*s*_) to the lower level, based on which the lower level would update its representation. Then, the higher-level prediction error (PE, *x*_*s*_) could be calculated as the difference between the top-down predictions and the upcoming signals. Next, *x*_*s*_ would propagate back to the higher level so as to optimize the next top-down prediction. At the lower level, its PE (*x*_*w*_) was simulated as cosine distances between predicted and real vectors of words, which is a robust index to reflect the dissimilarity and is less sensitive to the magnitude of vectors compared to other metrics (see Materials and Methods). In the continuous updating PC model, the prediction and PE were allowed to be transferred between the lower and higher levels instantaneously (Fig. 4a, left). While in the sparse updating PC model, the information between higher and lower levels is transmitted with a delay Δ*t* (Fig. 4a, right), which ensures that predictions and PEs were only exchanged across levels around the sentence boundaries, aligning with the previous findings [26,65–67]. Neural responses were simulated using these PC models and then transformed into the BOLD signals [68,69]. Finally, the gradient descent algorithm was applied to estimate the parameter of the PC models, and the mean square error (MSE) was calculated between the simulated BOLD signals and the real BOLD signals to index the performance of these PC models (Fig. 4b, c). The lower the MSE, the better the model’s performance (see Materials and Methods).

**Figure 4.**
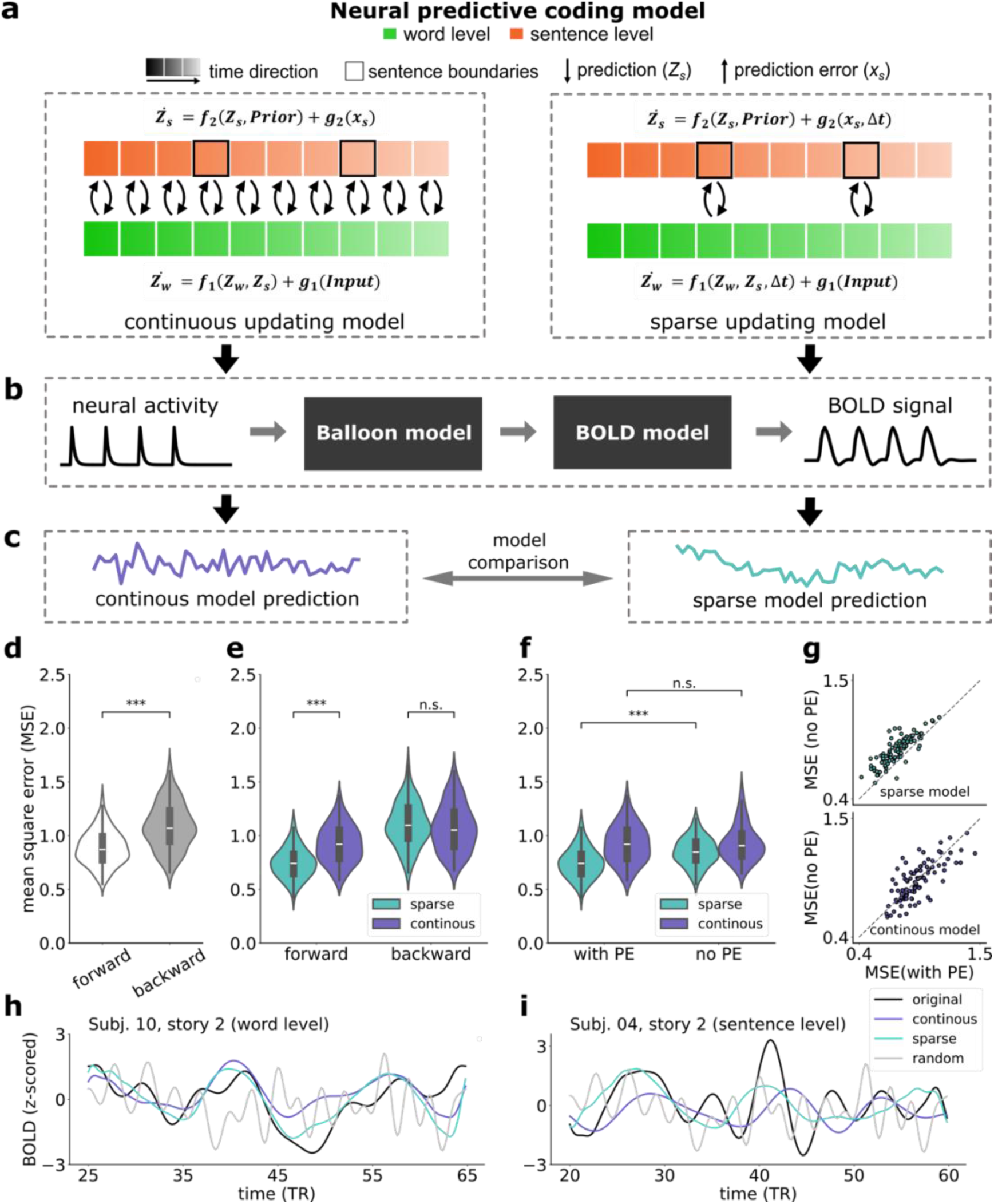
Results of predictive coding (PC) neural modeling. a) Two computational models were built according to the continuous updating hypothesis and the sparse updating hypothesis. Left panel: the continuous updating hypothesis assumes that the higher level is updated continuously as inputs change over time. Right panel: the sparse updating hypothesis assumes that the higher level only predicts and updates at its preferred timescales (i.e., the sentence boundary). b, c) The simulated data generated by the models were transformed into the putative BOLD signals via the hemodynamic model, and further compared with the real fMRI responses for the model comparison. d) Model performance in forward and backward conditions. e) The sparse PC model performs better than the continuous PC model in the forward condition only. f) Model performance when prediction error is not involved. g) Comparison of MSE values for sparse and continuous models by leveraging PE for all subjects across stories. h, i) Examples of simulated fMRI signals from different PC models at both word (i.e., Subject 10, story 2) and sentence levels (Subject 04, story 2) are presented along with the corresponding real BOLD signals for illustration. Significant levels are set as *p* ≤ 0.001 (***), *p* ≤ 0.01 (**), *p* ≤ 0.05 (*), and *p* > 0.05 (n.s.).

**Figure 5.**
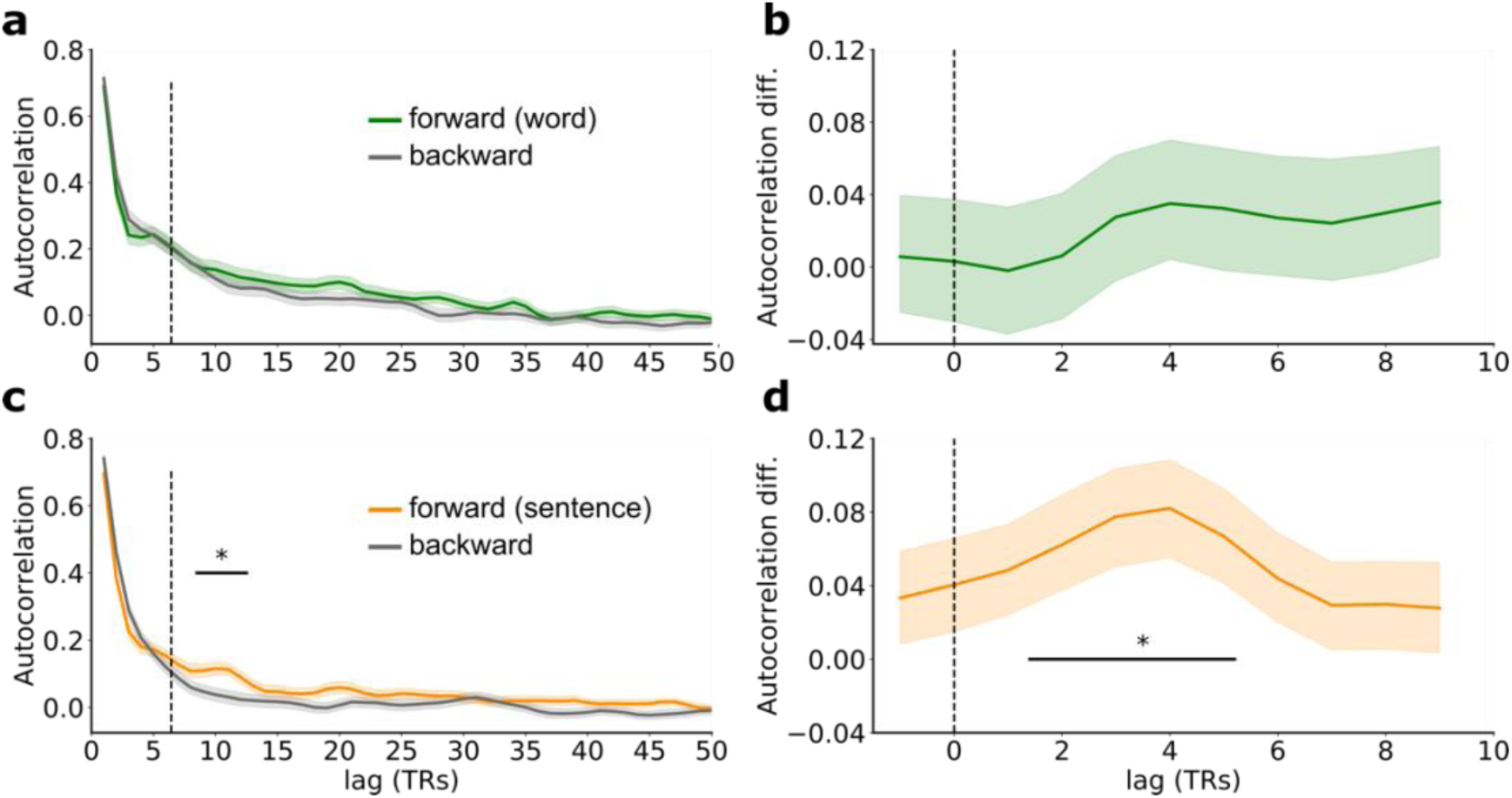
Results of the autocorrelation analysis. a, c) The autocorrelation results for the word and sentence levels respectively. The colored lines represent the forward condition; the gray lines represent the backward condition. b, d) The autocorrelation differences between forward and backward conditions were calculated and plotted for the selected windows at the sentence boundary. The vertical black dashed line represents the averaged sentence boundaries. The multiple corrections were controlled by the Bonferroni method with a threshold of *p* < 0.01.

First, MSE was compared between the forward and backward conditions. We found that both PC models performed better in the forward condition than the backward condition (paired *t*-test; *t*(185) = -12.760, *p* < 0.001, *d* = 1.086; Fig. 4d). However, the sparse PC model significantly outperformed the continuous PC model in the forward condition (paired *t*-test; *t*(92) = -17.438, *p* < 0.001, *d* = 1.110; Fig. 4e), but none was found in the backward condition (paired *t*-test; *t*(92) = 0.990, *p* = 0.325, *d* = 0.137; Fig. 4e). Additionally, this result was consistent when the analysis was conducted on each story separately (Fig. S6). The above findings suggested the sparse updating model could better explain the information update mechanism in the prediction hierarchy.

In addition, previous research has found that the lower level PE (*x*_*w*_) is crucial for linguistic processing [13] and plays an important role in demarcating events [70]. Thus, we tested this account by removing lower level PE (*x*_*w*_) from the models to validate the above results (i.e., when *x*_*w*_ was set to 0). The results confirmed this expectation with the sparse PC model (paired *t*-test; *t*(92) = -12.246, *p* < 0.001, *d* = 0.777; Fig. 4f, g), but not with the continuous PC model (paired *t*-test; *t*(92) = 0.787, *p* = 0.433, *d* = 0.060) in the forward condition (Fig. 4f, g).

Together, these computational modeling results directly supported the sparse updating hypothesis about the cortical architecture for hierarchical linguistic prediction.

### Validating the sparse updating model through an autocorrelation analysis

To further validate the sparse updating process, we additionally tested the autocorrelation effect in the BOLD signals. According to the sparse updating hypothesis, brain responses associated with the sentence prediction should not exhibit a significant change until the boundary of the sentence appears. In this sense, brain responses should be highly similar before the boundary of a sentence but should significantly differ across the boundary of a sentence. So, brain responses should exhibit a pattern of periodicity at the rate of sentence occurrence [26]. We tested this effect using the autocorrelation method on brain regions associated with the predictive representations of words and sentences. Specifically, we temporally shifted the BOLD signals without pre-whitening forward, with time lags ranging from 1 TR to 50 TRs, and correlations were calculated between the shifted signals and the original signal for each time lag. The autocorrelations were compared between the forward and backward conditions.

Our results showed a significantly stronger correlation, i.e., the autocorrelation effect, in the forward condition than in the backward condition in brain regions associated with sentence prediction, when the time lag was 8-11 TRs (Fig. 5c, d; Fig. S7). This time lag roughly time-locked to the length of sentences (3.43 TRs, Fig. S1c) with a hemodynamic delay-to-peak effect in BOLD signals (2.5 TRs, 5 seconds) (*ps* < 0.01, Bonferroni corrected. Fig. 5d). However, this effect was not found in the BOLD signals associated with the representation of word prediction (Fig. 5a, b). These findings provided additional support for the sparse updating hypothesis.

## Discussion

The present study aimed to test the hierarchical linguistic prediction on a longer timescale (i.e., sentence) than words and phonemes and understand the neurocomputational mechanisms underlying the interactions across different levels of the hierarchy. First, our results showed that the predictive representations of words are associated with brain responses in the STG and MTG, while those of sentences are associated with the TPJ, mPFC, and precuneus. Previous studies have consistently indicated that language comprehension inherently involves prediction, and linguistic prediction is hierarchically organized in the human brain [2–4,7]. However, most previous studies addressing linguistic prediction were conducted on phoneme and word levels. Some studies found that the predictions of phonemes were associated with the primary auditory cortex, while the predictions of words were associated with nearby associative brain regions such as the STG [1,12]. Although studies on language comprehension implied that the prediction of sentences might be related to high-level brain regions such as the TPJ and PFC, no studies have tested this possibility directly. By combining the NLP approach with the fMRI technique, here we not only confirmed the association of word prediction with the temporal cortex but revealed a novel association of sentence prediction with the core regions of the DMN (TPJ, mPFC, and precuneus).

There are studies, however, on word prediction whose results differed from ours. For instance, some studies found an association between word prediction with widely distributed brain regions in the lateral frontal cortex, in addition to the bilateral STG [1,13,14,71–74]. One possibility for this discrepancy is that these studies tapped into word prediction with lexical processing difficulties such as cloze probability [75] or entropy [76] rather than the predictive representation itself. We postulated that the involvement of the frontal cortex may result from the processing difficulty and the corresponding process of cognitive control [77,78]. However, Goldstein et al. (2022) employed encoding model and found that the IFG was also significantly involved in word prediction [13]. Although the authors ruled out the potential context effect, their control analysis was conducted for the averaged signal across all significant electrodes including both IFG and STG electrodes, leading to difficulties in disentangling the potential differences between IFG and STG. The present study investigated the neural underpinnings of linguistic prediction per se rather than processing difficulty and meanwhile controlled the potential confounding factors from the past linguistic context.

Thus, our results provided more direct evidence supporting the anatomical architecture for hierarchical linguistic prediction.

Furthermore, it has been proposed that DMN (especially for mPFC, precuneus, and TPJ) plays a key role in processing naturalistic stimuli [32], such as written or spoken stories [10,33] and movies [79]. To investigate its potential functions, on the one hand, some of the previous studies scrambled the real-life stories at different timescales (e.g., word, sentence, paragraph, etc.) [10,25] or shuffled parts of the stories to create different versions [32,80]. By comparing neural responses across different versions, researchers have found that the DMN is responsible for integrating external information with relatively long prior context (from seconds to minutes) [32,80,81]. On the other hand, another possible cognitive process associated with the DMN during narrative processing is using stored information to simulate possible future events and plan ahead (i.e., the prospective memory) [31,82]. For example, participants were asked to imagine a plausible event that had not happened before while lying in an fMRI scanner [31,83], and their neural signals in DMN regions such as the mPFC and precuneus showed significant activation. Interestingly, this “future envisioning” map largely overlaps with regions involved in episodic memory, hence facilitating the development of the constructive episodic simulation hypothesis [31]. These findings suggest that one of the important functions of DMN is to enable simulation of future events based on past events. This perspective is highly consistent with the concept of pre-activation in linguistic prediction [82,84]. While anticipatory signals in DMN regions have been extensively detected [29,30], the timescale underlies prospective prediction of the DMN is still not clear. In our study, we identified an involvement of the TPJ and DMN midline core areas in sentence prediction, providing novel evidence for the DMN’s role in predicting linguistic units with longer timescales.

The neural representations of the word and sentence prediction may be reminiscent of the research on the temporal receptive window (TRW), which concentrates on the representations of past linguistic contexts at multiple timescales [10,11,81]. These studies have proposed a temporal hierarchy of context in the cerebral cortex, stretching from the early sensory regions such as the superior temporal gyrus (STG) responding to shorter timescales (i.e., small TRW) to higher-level brain regions such as the temporoparietal junction (TPJ) and prefrontal cortex (PFC) responding to longer timescales (i.e., large TRW). These evidence suggests the existence of two types of temporal hierarchies in the brain [30], i.e., a prospective hierarchy that involving prediction at multiple timescales into the future, and a retrospective timescale hierarchy that representing contextual information from the past. However, the relationship between these two hierarchies remains unclear, posing an open question for further investigation.

Additionally, our study found a strong right lateralization of neural patterns for sentence prediction. While recent studies have challenged the traditional view on naturalistic language comprehension in the left hemisphere, showing instead that the process involves bilateral hemispheres [85–87], the underlying function of the right hemisphere in the prediction process remains unclear. Our results suggest that the default mode network (DMN) of the right hemisphere plays a dominant role in sentence prediction, shedding novel light on a potential function of the right brain. Furthermore, previous research on sequential music processing has emphasized the importance of the right hemisphere in perceptual segmentation and coarse-grained event boundaries in music [88], supporting the notion that the right hemisphere may have a distinct function in processing longer timescale information.

Second, our computational modeling results supported the sparse, rather than continuous, updating strategy during hierarchical linguistic prediction. Previous research that supported the continuous updating hypothesis usually employed correlation approaches such as inter-subject pattern correlation (ISPC) [25] or cross-context correlation [24]. The former examined brain response spatial similarities between subjects for each moment in time, while the latter calculated the neural similarities across different trials for each time point. These approaches, however, might mix up the effect of information updating and accumulating, thus may not be able to specifically disentangle different effects. Besides, although Chien & Honey (2020) also employed a computational modeling approach, they only tested one of the two rival hypotheses, leaving an open question of hypotheses comparison [25]. Most importantly, no studies have tested these two hypotheses at the sentence level. In the present study, we employed the computational modeling approach to uncover the hidden mental states (i.e., information updating at the sentence level) and compared two competing models to directly test the two rival hypotheses (i.e., the continuous and sparse updating hypotheses). The results supported the sparse updating hypothesis, which provides novel insight into the computational mechanisms of information updating in the hierarchy of linguistic prediction. Additionally, the sparse updating model has been proven to be more computationally efficient and resource-saving than the continuous updating model [35]. Thus, our findings also provide empirical support for the brain’s economy principle; that is, the human brain is organized to carefully manage the input resources in the services of delivering robust and efficient performance [89,90].

However, recent eye-tracking and electroencephalogram (EEG) studies provide support for the incremental nature of language processing [75,91,92], which seemingly aligns more with the continuous updating strategy [93]. For instance, some EEG studies used the rapid serial visual presentation (RSVP) paradigm where each word was presented one at a time in the center of the screen. They found that the accruing sentence contexts affect the N400 activity for each word [75,93]. Leon-Cabrera et al (2019) also presented participants with sentences via RSVP paradigm and observed a slow, sustained negative drift throughout the sentence processing, which could potentially reflect the continuous construction of the meaning representation [94]. However, we believe that the sparse updating strategy does not necessarily contradict to, but could be accommodated with the incremental online language processing. In our model, the sparsely updated information over two levels does not influence the bottom-up input (i.e., the word level prediction error *x*_*w*_(*t*)), allowing the lexical information to be accumulated instantaneously at the word level. Moreover, updating strategies might be different between levels of the prediction hierarchy. As only the interaction between the word and sentence levels have been investigated in the present study, more future research is required to further elucidate this important question.

There were several limitations of the present study. First, we cannot obtain the real- time attentive states of participants because either regular or random assessments during story listening would disrupt the processing of continuous speech and the real-time predictions using the preceding contexts. Second, although fMRI has a high spatial resolution, its temporal resolution is relatively low to precisely describe the dynamic process of linguistic prediction on a smaller timescale such as phonemes. Moreover, we did not analyze the subcortical regions. The contribution of subcortical regions such as the cerebellum might be overlooked. We take our findings as a first step in this novel direction and hope more studies will be conducted in the future.

In conclusion, by directly examining the predictive representations of linguistic units in the brain, we revealed that the predictive representations of linguistic units with different temporal sizes showed a gradient architecture from the temporal cortices to the DMN regions. Most importantly, our results revealed the underlying neural computational mechanisms of interaction between levels of the linguistic prediction hierarchy, supporting the sparse updating hypothesis. Together, these findings extend the understanding of the cortical architecture for hierarchical linguistic prediction and provide novel insight into the neurocomputational mechanisms of linguistic prediction hierarchy during narrative comprehension.

## Materials and Methods

### Participants

Neuropower (http://neuropowertools.org/) was used to determine the sample size before the experiment (28 participants were recommended when statistical power was 0.8). Totally 38 healthy native Chinese speakers participated in this study. All participants were right-handed [95] and without hearing, psychiatric, or neurological problems according to their self-report. Six participants were removed because of excessive head motions (larger than 3 mm or 3 degrees) and one was removed because he fell asleep during the task, leaving thirty-one participants (mean age: 23 years, ranging from 19 to 26; 19 females) with valid data.

The study protocol was approved by the Institutional Review Board of the State Key Laboratory of Cognitive Neuroscience and Learning at Beijing Normal University. Written informed consent was obtained from all participants.

### Stimuli

For the narrative listening task, it is routine to have multiple runs in a study [14,49,79], with the purpose of including more stimuli to increase the reliability of statistical tests as well as to reset the fatigue state of participants and reduce technical problems (for example, scanner overheating). Therefore, three stories were included in this study, which could provide us with higher inference power when applying our conclusions to the natural contexts in daily life. Specifically, story 1 and 2 were obtained by asking two female speakers to freely introduce “*an unforgettable experience in your college life*”, while story 3 was recorded by asking a female speaker to read a text adapted from *The Kite Runner*. The stories were recorded by the audio system of FOMRI III (Optoacoustics Ltd.) and were further denoised using Audacity (https://www.audacityteam.org/). These stories were matched regarding their perceptual features such as clarity, familiarity, and complexity (see Task and Procedures below). Additionally, each audio was temporally inversed to be used in the backward condition to control for acoustic features.

### Task and procedures

Before the experiment, an appropriate sound volume was set to a comfort level according to each participant’s subjective report. During the experiment, participants were requested to passively listen to the three stories (the forward condition) and the corresponding control audios (i.e., the temporally inversed audio, the backward condition) while fMRI data were collected. During story listening, participants were also asked to fixate on a cross at the center of the black screen simultaneously. The sequence of the six audios (three in the forward and three in the backward conditions) was counterbalanced across participants. An interval was inserted between two audios during which the participants could take a rest. The length of the interval was flexible and determined by the participants. Additionally, all audios were preceded by 10 s silence with a black screen to control T1 equilibration effects [14,30]. Audios were played with OptoACTIVE headset, which can actively eliminate the noise produced by the MRI scanner instantaneously. This device has been widely used in previous auditory studies [96–98]. E-prime (v2.0.10) software (https://pstnet.com/products/e-prime/) was used to control stimuli presentation.

At the end of each story, the participants were tested on how well they perceived and comprehended the story. Specifically, at the perceptual level, the participants were requested to judge clarity, familiarity, and complexity on a 5-point Likert scale (1 was the lowest level, and 5 was the highest level). At the comprehension level, several True or False questions were designed for each story based on the richness of contents (3 questions for story 1 and 2; 5 questions for story 3). These questions were either about the details (mentioned only once in a story) or the gist (mentioned several times in a story) [99]. The questions of each story covered both types of questions. Statistical tests were conducted on both scores to determine how well the participants perceived and comprehended these three stories.

Additionally, to validate these questions, additional 21 participants who were not included in the formal experiment and were naïve to the purpose of the experiment were recruited. They were asked to judge “How well do you think these questions could reflect the listener’s comprehension of the story?” on a 7-point Likert scale (1 was strongly disagree, and 7 was strongly agree). One sample *t*-test was performed on the scores against the uncertain level (i.e., 3.5 on the 7-point scale). The FDR method was used to correct for multiple comparisons [52]. The results showed that the scores of all 3 stories were significantly larger than the uncertain level (story 1: 5.71 ± 0.78, *t*(20) = 10.02, *p* < 0.05; story 2: 5.43 ± 1.08, *t*(20) = 6.09, *p* < 0.05; story 3: 5.81 ± 0.80, *t*(20) = 9.60, *p* < 0.05), indicating that these questions were reliable in reflecting the level of comprehension on the stories used in this study.

### Statistical analysis

For both behavioral and neural data, the Agostino D test was used to determine the normality of data distribution before conducting statistical analysis. If data were subject to the normal distribution, a parametric test would be used (e.g., paired *t*-test); otherwise, the non-parametric test would be used (e.g., Wilcoxon sign rank test). All statistical tests without further explanations were two-tailed with a threshold of *p* < 0.05.

### fMRI data acquisition and preprocessing

The fMRI data were acquired with Siemens TRIO 3-Tesla scanner at the Imaging Center for Brain Research, Beijing Normal University. The functional images were collected using an EPI sequence (TR = 2000 ms, TE = 30 ms, flip angle = 90°, FOV= 200 mm, voxel size = 3.1 × 3.1 × 3.5 mm^3^, interleaved). The structural T1-weighted images were collected using magnetization-prepared rapid gradient-echo (MP-RAGE) sequence (TR = 2530ms, TE = 3.39 ms, flip angle = 7°, FOV = 256 mm, 144 sagittal slices, voxel size = 1.3 × 1.0 × 1.3 mm).

DPABI toolbox built on SPM12 (www.fil.ion.ucl.ac.uk/spm/) was used for data preprocessing [100]. After removing the first 5 volumes corresponding to the silent period (10s), the images were slice-time corrected, spatially realigned to the first image in a run using the rigid body registration, and co-registered to their corresponding anatomical images. Next, both functional and anatomical images were normalized to the standard Montreal Neurological Institute (MNI) space, with functional images resampled to 2×2×2 mm^3^ voxel size. Then, data were spatially smoothed using Gaussian kernel with a 6 mm full width at half maximum (FWHM). Finally, all data were further detrended and temporally high-pass filtered (128 s cutoff), and nuisance regressed (including Friston’s 24 motion parameters and five principal components of the white matter and cerebrospinal fluid) [101].

### Obtaining the predictive representations of words and sentences

#### Dataset generation

Chinese Wikipedia derived from the project *Large Scale Chinese Corpus for NLP* (https://github.com/brightmart/nlp_chinese_corpus) was used as the corpus. First, during preprocessing, symbols and tokens that are unrelated to contents were removed to reserve most of the linguistic information embedded in the corpus. Then, the corpus was segmented into word units using jieba toolbox (https://github.com/fxsjy/jieba) and parsed into sentences with the end-of-sentence punctuations (i.e., period, question mark, exclamation mark, and ellipsis), respectively. Second, a document was randomly selected from the corpus, from which a linguistic unit (a word or a sentence) was randomly selected as the to-be-predicted target. All the texts prior to the target were assigned as the prior linguistic context. A dataset was generated for words and another one for sentences with this procedure, and each dataset included about 0.2 million items, with each item contained a target and its linguistic context.

We did not remove any functional words. Instinctively, the removal of functional words could eliminate some informative messages from the text, allowing NLP algorithms to be more focused on the semantics of content words. However, this approach may overlook the fact that the functional words, such as the negation words like “not”, “nor”, “never”, etc., also contain some semantics and particularly crucial syntactic information, which is particularly important for understanding the natural language. In fact, there is a debate about whether functional words should be removed when applying the BERT models. The original BERT model, for example, did not suggest removing any stop words [37]. Moreover, Qiao et al., (2019) found that removing functional words does not affect BERT model performance [102]. Additionally, Alzahrani & Jololian, (2021) showed that removing functional words even negatively affects the performances of the BERT model in a gender classification task [103], decreasing the classification accuracy from 86.67% to 78.86%. Therefore, to incorporate both semantic and syntactic information into the vector representations, we have decided to stay close to the original usage of the BERT model by including both functional and content words in the analysis to more focus on the intrinsic timescales rather than only the functional time scales in this study.

#### Vector representations

Both the prior linguistic context and the prediction target were vectorized using WWM- RoBERTa model [36]. WWM-RoBERTa is a type of the Bidirectional Encoder Representations from Transformers (BERT) language model [37]. BERT is a pre- trained representation model designed with a multi-layer bidirectional Transformer encoder conditioning on both left and right contexts [37]. The overarching mechanism of the transformer is multi-head self-attention, which is fundamentally the weighted mean of all the input vectors [104]. Compared with the original BERT model, WWM- RoBERTa model has a bigger architecture (the number of layers = 24, the hidden size = 1024, the number of self-attention heads = 16, total parameters = 340 M), with larger batch size and more dataset. More importantly, it is trained based on the prediction task of whole words rather than characters, hence possessing high generalizability and adaptability for Mandarin [38].

WWM-RoBERTa model was applied using python (v3.7) with bert-as-service module (https://github.com/hanxiao/bert-as-service). This module was developed to map a variable-length sentence to a fixed-length vector (1024 dimensions). Note that “sentence” here means a span of text from the corpus, usually much longer than a single sentence [37]. To obtain a comprehensive text vector, the bert-as-service averaged the output vector of the penultimate hidden layer of all the tokens in an input text, because the last layer representation is sensitive to the model training tasks (i.e., masked language model and the next sentence prediction). Note that there is also an alternative way to have the vector, i.e., to use the [CLS] token. [CLS] is a special symbol added in front of sentence inputs, which was frequently used to represent the overall information of the inputs [37]. However, previous studies have indicated that the effect produced by [CLS] embedding approach is weaker than that of the averaging approach [39,105,106]. Therefore, bert-as-service default setting, i.e., the average approach, was used in the present study. Specifically, for word units, we obtained vectorial representations where the target word is the only input to the BERT model (without contexts). For sentence and context information, we computed the averaged embedding of all words in the text. In addition, because of the quadratic relationship between the length of text sequence and computational cost [37], the maximum input length of WWM-RoBERTa model is 512 characters given a text. In this study, two additional separation tokens were inserted at the front ([CLS]) and the end ([SEP]) of the input text. Thus, the actual length of input was 510. Here a “split-and-average” method was used to cope with the restriction of the WWM-RoBERTa model on the length of input text (i.e., < 510 characters). That was, if the context length was larger than 510 but less than 1020, the context would be split into 2 parts equally, and the vectors of them were averaged. The averaged vectors were used as the vectorial representation of this context. The “split- and-average” method should be taken as an extension of the averaging approach. The same procedure was generalized to a longer context with a maximum length of 4080 (8 segmentations). Finally, for each item of the dataset, there were 1024-dimension vectors for the prior linguistic context and 1024-dimension vectors for the prediction target (a word or a sentence).

To validate the split-and-average method, we selected 1000 documents whose text had a length ranging from 50 to 510 randomly. First, the texts were converted into vectors using WWM-RoBERTa model to index the Whole Text Vector (WTV). Additionally, these texts were split into N segments (here *N* ∈ {2, 3, 4, …,8}). These segments were also converted into vectors and averaged to index the Segment Text Vector (STV). Second, cosine distances between the WTV and STV were calculated as Dorig. Third, the WTVs and STVs were randomly paired 1000 times. For each time, cosine distances between the WTV and the randomly paired STV were calculated as Drand. This procedure generated a null distribution, based on which *p*-values were obtained for Dorig. The result showed that Dorig were significantly larger than null distributions in all segment conditions (*ps* < 0.001, FDR corrected), indicating that the split-and-average method is valid.

#### Model building

The multiple ridge regression modeling method was used to delineate the prediction relationship between the prior linguistic contexts and the prediction targets. The model is composed of 1024 ridge regressions, with each dimension of the upcoming input vectors (1024 dimensions in total) being predicted using one ridge regression model based on the corresponding dimension of the linguistic context vectors. For the present, we understood very little about the meaning of each dimension outputted from the LLM. But a recent study by Huang et al., (2021) shows that a whitening-based vector normalization strategy, i.e., removing the covariance structure in the vectors, consistently enhances model performance [105]. Our method was inherently aligned with this concept. Mathematically, for each ridge regression model, given n samples of one dimension of the upcoming input vector ***Y*** (n × 1) and context matrix ***X*** (n × 1024), we expected to estimate coefficients *β* (1024 × 1) and *β*_0_ (n × 1) by minimizing the following cost function:

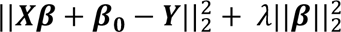

where *λ* is the regularization term for preventing model overfitting by reducing coefficients *β*.

To estimate the parameters (*β* and *β*_0_), the dataset (see “Dataset generation”) was further divided into the training set (80%) and the test set (20%) (Fig. 1a). The reasonable *λ* was evaluated using 4-fold cross validation within the training set. The input vector ***Y*** and the context matrix ***X*** were normalized by column (i.e., across training samples) in advance. The model training and testing processes were implemented with the sklearn toolbox [107].

#### Model validation

A pairwise classification task was used to evaluate the performance of the model [45]. To this end, first, 1000 samples were randomly selected from the testing set, and each sample contains a vector *Vreal-target* for the prediction target and a vector *Vreal-context* for the prior linguistic context. Second, a simulated vector *Vpred-target* for the prediction target was generated using the models trained above based on *Vreal-context*. The cosine distance between *Vpred-target* and *Vreal-target* was calculated as D1. Third, cosine distance D2 was calculated between *Vpred-target* and a randomly selected *Vrand-target.* If D1 was larger than D2, a label of “right” was assigned; otherwise, a “wrong” label was assigned. Additionally, we also applied the Pearson correlation index between the actual and predicted targets to supplement our classification results. Based on this procedure, a percentage of accuracy could be obtained. Finally, these procedures were repeated 1000 times for robustness.

### Transferring the model to the current stimuli

The same procedures as mentioned above were applied to the experimental materials used in the present study. Briefly, the texts of the three stories were segmented at both word and sentence levels (Fig. 1b). Word was separated using jieba toolbox based on Python. To partition sentences appropriately, we recruited 10 raters to segment the text wherever they thought there should be the end of a sentence (sentence boundaries annotation task). A sentence boundary was made if at least 5 raters made marks [26]. Then, a trained experimenter further examined and refined the positions of marks. Additionally, Praat (http://www.praat.org/) was utilized to align the segmented text (words and sentences) to audios. These aligned data were further transformed into vectorial representations.

### Relating the predictive representations of linguistics units with BOLD signals using gGLM analysis

A gGLM analysis was performed to identify the neural underpinnings of the predictive representations of words and sentences respectively. A leave-one-subject-out cross validation (LOSO-CV) approach was applied, where the model performance of a specific participant was evaluated based on the data of all other participants. This approach could effectively avoid overfitting and suppress the non-independence error [51]. Additionally, Because the dimensions of the vectorial representations are relatively large (i.e., 1024 dimension), which might cause the overfitting problem during gGLM analysis, a feature reduction approach was utilized by combining Isomap and PCA methods [108,109]. Previous research has shown that the concatenated components of Isomap and PCA could have a comparable performance with the original data [108]. To meet the minimum criteria (i.e, PCA cumulative variances explained >= 50% and Isomap residual variance reaches the least value), 15 components were selected from Isomap and 35 principal components were selected from PCA (Table S2, S3).

Furthermore, the design matrices were generated using the function ‘make_first_level_design_matrix’ within the python toolbox nilearn (https://nilearn.github.io; for more details, please refer to https://nilearn.github.io/dev/modules/generated/nilearn.glm.first_level.make_first_lev el_design_matrix.html). Specifically, the following steps were conducted in the function (Fig. S2): 1) Oversampling. Based on the timing information of each language unit in the stories (such as the offsets of each word or sentence), a time course was generated and then oversampled to a sampling rate of 50Hz; 2) HRF convolution. The oversampled time courses were convolved with HRF; 3) Downsampling. The convolved time course was further downsampled to 0.5Hz, which corresponded to the sampling rate of fMRI (i.e., TR = 2 s). The downsampled time course were used to fit the fMRI time course using encoding models.

The variance partitioning (VP) approach was employed to identify the prediction effects. Specifically, 1) a model only including the context representations (MC) was used to obtain the context-specific effect, the context representations were obtained with the vectorial representations of prior linguistic context generated with the same procedures mentioned above (Table S2, S3); 2) a full model including both the prior linguistic context and the predictive representation (MF) was used to obtain both context and predictive effects. The explained variance (*R^2^*) was compared between MF and MC, and the difference (i.e., MF - MC) is defined as the actual effect of predictive representation. In addition, following the concept of pre-activation [3], the predictive representations of linguistic unit N (i.e., word/sentence) were aligned to the offset of linguistic unit N-1 (Fig. S2). For each model, the identity of each participant and each story was dummy coded and used as covariates to control for the individual differences and stories’ differences. The regressors modeling the boundaries of each word/sentence were included to account for the temporal delay in BOLD activation respective to stimulus presentation. Additionally, the log-transformed word frequencies or sentence frequencies (i.e., the average frequency of all words in this sentence) were also regressed out from the corresponding model to control for the statistical influence of everyday usage (Willems et al., 2016). Word frequencies were obtained from Cai & Brysbaert (2010), which is derived from the film subtitles that can capture the variability of the word’s daily usage [111]. The frequency was log-transformed because of its inherent skewed distribution.

The gGLM analysis was performed in a parcellation approach, where 400 non- overlapping parcels were used [46]. The BOLD signals within each parcel were pre- whitened with the AR(1) noise model implemented in nistats [112] and averaged. The same procedures were applied to the forward and backward conditions. Paired t-test between the forward and backward conditions was performed to select significant parcels. Multiple comparison problem was solved by a threshold determined by FDR (*q* < 0.01) [52]. The significant results were projected to a cortical surface for visualization using Brainnet Viewer [113].

#### Permutation test

To validate the neural underpinnings of predictive representation, a permutation test was performed. Specifically, the rows and columns of the design matrices were shuffled 1000 times to remove the prediction effect. Then, the same pipeline as mentioned above was repeated to generate a null distribution of *R^2^*. Finally, *p* values were obtained by calculating the position of the original *R^2^* value in this null distribution.

### Computational modeling

#### Building models

To directly test the sparse and continuous updating hypotheses, two computational models that meet the minimal assumptions of the predictive coding framework [56,114] were built. The PC framework postulates that the brain hierarchically processes upcoming information, where level N generates prediction signals to level N-1. Then, the prediction errors (PE, i.e., the difference between the neural responses at level N and the upcoming neural responses at level N-1) are sent back to update the next prediction at the N level [57]. The PC framework has been theoretically illustrated and tested with BOLD signals in many studies [14,57,62,114–116]. In our models, the architecture aligned with the continuous updating hypothesis can be described with differential equations below (continuous updating PC models):

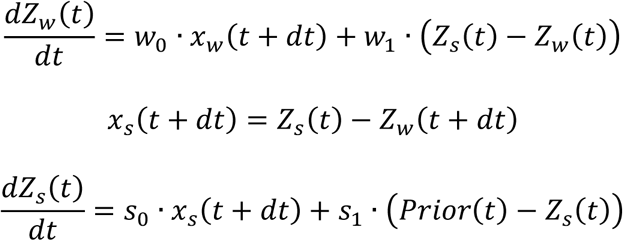

where *dt* was set to be the repetition time (TR = 2s) during model estimation. *x*_*w*_(*t*) is the PE at the word level, which is substituted by the min-max normalized cosine distances between predicted vectors and real vectors of words, and resampled to fMRI acquisition rate (TR = 2s). Cosine distance between linguistic vectors was used to calculate the errors in the present study because it is a robust index to reflect the dissimilarity and is less sensitive to the magnitude of vectors compared to other metrics [117–119]. Furthermore, the PEs in our model were restricted to the positive value, so we implemented the min-max normalization on the cosine distance. *Z*_*w*_ and *Z*_*s*_ are the BOLD signals associated with the predictions of words and sentences respectively, which are the averaged BOLD signals across the parcels (i.e., *Z*_*w*_ is the average of bilateral STC and MTC; *Z*_*s*_ is the average of the right TPJ, medial PFC and precuneus).

The brain regions were identified in the gGLM analyses. The *x*_*s*_(*t*) are the PE at the sentence level, which is modeled here as the difference between *Z*_*s*_(*t*) and the upcoming *Z*_*w*_(*t*). *Prior*(*t*) is assumed to arise from the higher level of the hierarchy and exert a top-down influence on the neural responses at the sentence level *Z*_*s*_(*t*).

*Prior*(*t*) here was simply set as 0 in line with the previous research, assuming that the top-down prior has minimal influences on the information updating strategy at the current levels [56]. In addition, *w*_0_, *w*_1_, *s*_0_, and *s*_1_ are parameters needed to be estimated. Mathematically, *w*_0_ and *s*_0_ controls the strength of how PEs at the word and sentence levels affect neural signals, while *w*_1_ and *s*_1_ primarily regulate the decay rate of the *Z*_*w*_(*t*) and *Z*_*s*_(*t*), respectively.

By contrast, the sparse updating hypothesis suggests a discretized information exchange between adjacent levels. The model that aligns with this hypothesis can be described by the below differential equations with delays (sparse updating PC models):

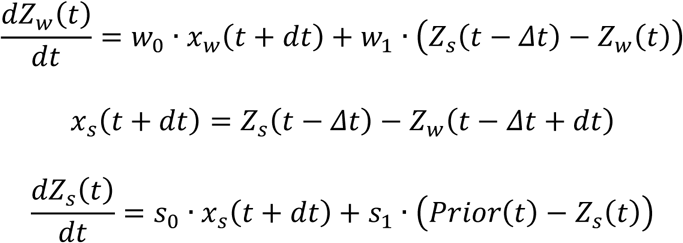

where *Δt* is a varying parameter quantifying the time lag between the current time point and the boundary of the last sentence. This parameter allows us to model the information updating process at the sentence boundaries rather than at every time point when a word occurs. All the other variables and parameters are the same as those in the continuous updating PC model.

To compare with the simulated and real fMRI signals, the neural signals simulated by different PC models were transformed into BOLD responses through the hemodynamic model, containing the Balloon model and the BOLD model [68,69,120]. Specifically, the Balloon model describes how neural responses cause changes in blood volume and deoxy-hemoglobin (dHb), and can be formulated as follows:

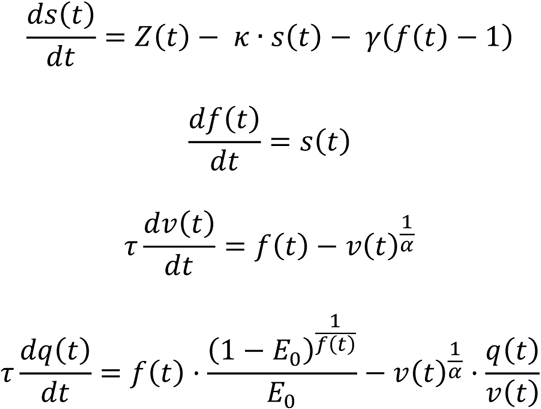

where *Z*(*t*) is the neural responses derived from the aforementioned PC models; *s*(*t*) represents vasodilatory signal; *f*(*t*) is the blood flow; *v*(*t*) is the local change in the blood volume in the blood vessel; *q*(*t*) represents the proportion of dHb.

Furthermore, the BOLD model characterizes how blood volume and dHb synergistically lead to changes in BOLD signal, which can be described by the following non-linear equation:

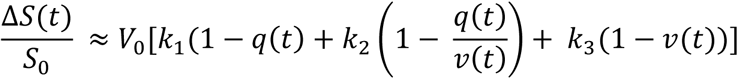

The parameters *k*_1_, *k*_2_, and *k*_3_ can be calculated through the following equations:

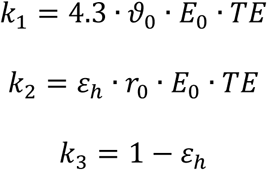

in which *S*_0_ is the BOLD signal at rest, and Δ*S* is the BOLD signal changes due to task performance. The detailed meanings of all the parameters in the hemodynamic and BOLD models are listed in Table S4. The simulation data for all hidden variables are visualized in Fig. S8.

#### Estimating models

The gradient descent method was employed to estimate parameters of *w*_0_, *w*_1_, *s*_0_, and *s*_1_. Gradient descent is an optimal algorithm to iteratively find the local minimum. Specifically, these four parameters (*θ* = {*w*_0_, *w*_1_, *s*_0_, *s*_1_}) are simultaneously updated:

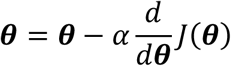

where *α* is the learning rate. Then the cost function *J*(*θ*) was defined as:

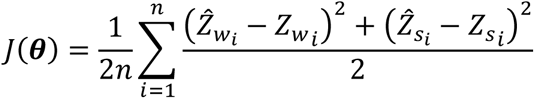

where *Z*_*w*_ and *Z*_*s*_ s sorememtiortm, are the fMRI signals associated with the predictions of words and sentences, which are the result of gGLM analysis for each level, before averaging and zscoring; *Z*^_*w*_ and *Ẑ*_*s*_ are the corresponding estimated signals; n is the length of neural signals in TR. During the model training process, learning rate *α* was set as 1 × 10^−5^, the convergence threshold was defined as the change of cost function *dJ* < 1 × 10^−4^. Because the cost function cannot be proven as concave, we randomized parameters *θ* for 10000 times to find the best initial condition. We applied the leave-one-subject-out cross validation approach to estimate *J*(*θ*) . The resulting mean squared error (MSE), which is two times of *J*(*θ*), was reported in this study to index the performance of the models.

### Autocorrelation analysis

The BOLD signals in the parcels that survived the statistical test were used to calculate the autocorrelation effect. The time courses of the signals were temporally shifted forward from 1 TR to 50 TRs. Then, Pearson correlation was calculated between the original signals and the shifted signals within each participant, using tsa.acf() function from statsmodels toolbox [121].

## Data Availability

The dataset will become available upon request.

## Code Availability

All the scripts for analyses will become available upon request.

## Acknowledgements

We thank Xiangyu He, Xiaofang Lu, and Xinran Xu for helping collect the fMRI data, Amirhossein Khalilian-Gourtani for the advice and assistance on the neural modeling, and other members of Lu lab and Flinker lab for extensive discussions. This work was supported by the National Natural Science Foundation of China (62293550, 62293551, 61977008).

## Author contributions

C.L. and F.Z. conceived the project; A.F., S.Z. and Y.L. contributed ideas for experiments and analysis; F.Z. collected data and performed the analyses; C.L. and A.F. critically revised the article; C.L. and F.Z. finished the manuscript with input from all authors.

## References

1. Heilbron M, Armeni K, Schoffelen J-M, Hagoort P, de Lange FP. A hierarchy of linguistic predictions during natural language comprehension. Proc Natl Acad Sci U S A. 2022;119: e2201968119. doi:10.1073/pnas.2201968119

2. Kuperberg GR, Jaeger TF. What do we mean by prediction in language comprehension? Lang Cogn Neurosci. 2016;31: 32–59. doi:10.1080/23273798.2015.1102299

3. Pickering MJ, Gambi C. Predicting While Comprehending Language: A Theory and Review. Psychol Bull. 2018;144: 1002–1044. doi:10.1037/bul0000158

4. Pickering MJ, Garrod S. Do people use language production to make predictions during comprehension? Trends Cogn Sci. 2007;11: 105–110. doi:10.1016/j.tics.2006.12.002

5. Forseth KJ, Hickok G, Rollo PS, Tandon N. Language prediction mechanisms in human auditory cortex. Nat Commun. 2020;11: 14. doi:10.1038/s41467-020-19010-6

6. Hagoort P. The neurobiology of language beyond single-word processing. Science. 2019;366: 55-+. doi:10.1126/science.aax0289

7. Lewis AG, Bastiaansen M. A predictive coding framework for rapid neural dynamics during sentence-level language comprehension. Cortex. 2015;68: 155–168. doi:10.1016/j.cortex.2015.02.014

8. Bornkessel-Schlesewsky I, Schlesewsky M, Small SL, Rauschecker JP. Neurobiological roots of language in primate audition: common computational properties. Trends Cogn Sci. 2015;19: 142–150. doi:10.1016/j.tics.2014.12.008

9. Ding N, Melloni L, Zhang H, Tian X, Poeppel D. Cortical tracking of hierarchical linguistic structures in connected speech. Nat Neurosci. 2016;19: 158-+. doi:10.1038/nn.4186

10. Lerner Y, Honey CJ, Silbert LJ, Hasson U. Topographic Mapping of a Hierarchy of Temporal Receptive Windows Using a Narrated Story. J Neurosci. 2011;31: 2906–2915. doi:10.1523/jneurosci.3684-10.2011

11. Yeshurun Y, Nguyen M, Hasson U. Amplification of local changes along the timescale processing hierarchy. Proc Natl Acad Sci U S A. 2017;114: 9475–9480. doi:10.1073/pnas.1701652114

12. Donhauser PW, Baillet S. Two Distinct Neural Timescales for Predictive Speech Processing. Neuron. 2020;105: 385-+. doi:10.1016/j.neuron.2019.10.019

13. Goldstein A, Zada Z, Buchnik E, Schain M, Price A, Aubrey B, et al. Shared computational principles for language processing in humans and deep language models. Nat Neurosci. 2022;25: 369–380. doi:10.1038/s41593-022-01026-4

14. Willems RM, Frank SL, Nijhof AD, Hagoort P, van den Bosch A. Prediction During Natural Language Comprehension. Cereb Cortex. 2016;26: 2506–2516. doi:10.1093/cercor/bhv075

15. Fedorenko E, Scott TL, Brunner P, Coon WG, Pritchett B, Schalk G, et al. Neural correlate of the construction of sentence meaning. Proc Natl Acad Sci. 2016;113: E6256–E6262. doi:10.1073/pnas.1612132113

16. Woolnough O, Donos C, Murphy E, Rollo PS, Roccaforte ZJ, Dehaene S, et al. Spatiotemporally distributed frontotemporal networks for sentence reading. Proc Natl Acad Sci. 2023;120: e2300252120. doi:10.1073/pnas.2300252120

17. Hagoort P, van Berkum J. Beyond the sentence given. Philos Trans R Soc B-Biol Sci. 2007;362: 801–811. doi:10.1098/rstb.2007.2089

18. Meeds R, Bradley SD. The Role of the Sentence and Its Importance in Marketing Communications. Psycholinguistic phenomena in marketing communications. Mahwah, NJ, US: Lawrence Erlbaum Associates Publishers; 2007. pp. 103– 120.

19. Mellem MS, Jasmin KM, Peng C, Martin A. Sentence processing in anterior superior temporal cortex shows a social-emotional bias. Neuropsychologia. 2016;89: 217–224. doi:10.1016/j.neuropsychologia.2016.06.019

20. Gao P, Jiang Z, Yang Y, Zheng Y, Feng G, Li X. Temporal neural dynamics of understanding communicative intentions from speech prosody. NeuroImage. 2024;299: 120830. doi:10.1016/j.neuroimage.2024.120830

21. Yu S, Gu C, Huang K, Li P. Predicting the next sentence (not word) in large language models: What model-brain alignment tells us about discourse comprehension. Sci Adv. 2024;10: eadn7744. doi:10.1126/sciadv.adn7744

22. Shi W, Demberg V. Next Sentence Prediction helps Implicit Discourse Relation Classification within and across Domains. Proceedings of the 2019 Conference on Empirical Methods in Natural Language Processing and the 9th International Joint Conference on Natural Language Processing (EMNLP-IJCNLP). Hong Kong, China: Association for Computational Linguistics; 2019. pp. 5789–5795. doi:10.18653/v1/D19-1586

23. Schmitt L-M, Erb J, Tune S, Rysop AU, Hartwigsen G, Obleser J. Predicting speech from a cortical hierarchy of event-based time scales. Sci Adv. 2021;7: eabi6070. doi:10.1126/sciadv.abi6070

24. Norman-Haignere SV, Long LK, Devinsky O, Doyle W, Irobunda I, Merricks EM, et al. Multiscale temporal integration organizes hierarchical computation in human auditory cortex. Nat Hum Behav. 2022;6: 455–469. doi:10.1038/s41562-021-01261-y

25. Chien HYS, Honey CJ. Constructing and Forgetting Temporal Context in the Human Cerebral Cortex. Neuron. 2020;106: 675-+. doi:10.1016/j.neuron.2020.02.013

26. Baldassano C, Chen J, Zadbood A, Pillow JW, Hasson U, Norman KA. Discovering Event Structure in Continuous Narrative Perception and Memory. Neuron. 2017;95: 709-+. doi:10.1016/j.neuron.2017.06.041

27. Rubio-Fernandez P, Jara-Ettinger J. Incrementality and efficiency shape pragmatics across languages. Proc Natl Acad Sci U S A. 2020;117: 13399–13404. doi:10.1073/pnas.1922067117

28. Manning CD, Clark K, Hewitt J, Khandelwal U, Levy O. Emergent linguistic structure in artificial neural networks trained by self-supervision. Proc Natl Acad Sci. 2020;117: 30046–30054. doi:10.1073/pnas.1907367117

29. Bar M. The proactive brain: using analogies and associations to generate predictions. Trends Cogn Sci. 2007;11: 280–289. doi:10.1016/j.tics.2007.05.005

30. Lee CS, Aly M, Baldassano C. Anticipation of temporally structured events in the brain. Elife. 2021;10. doi:10.7554/eLife.64972

31. Schacter DL, Addis DR, Buckner RL. Remembering the past to imagine the future: the prospective brain. Nat Rev Neurosci. 2007;8: 657–661. doi:10.1038/nrn2213

32. Yeshurun Y, Nguyen M, Hasson U. The default mode network: where the idiosyncratic self meets the shared social world. Nat Rev Neurosci. 2021. doi:10.1038/s41583-020-00420-w

33. Xu J, Kemeny S, Park G, Frattali C, Braun A. Language in context: emergent features of word, sentence, and narrative comprehension. Neuroimage. 2005;25: 1002–1015. doi:10.1016/j.neuroimage.2004.12.013

34. Yarkoni T, Speer NK, Zacks JM. Neural substrates of narrative comprehension and memory. Neuroimage. 2008;41: 1408–1425. doi:10.1016/j.neuroimage.2008.03.062

35. Chung J, Ahn S, Bengio Y. Hierarchical Multiscale Recurrent Neural Networks. ArXiv. 2017;abs/1609.01704.

36. Cui Y, Che W, Liu T, Qin B, Yang Z, Wang S, et al. Pre-Training with Whole Word Masking for Chinese BERT. 2019; arXiv:1906.08101.

37. Devlin J, Chang M-W, Lee K, Toutanova K. BERT: Pre-training of Deep Bidirectional Transformers for Language Understanding. Minneapolis, Minnesota: Association for Computational Linguistics; 2019. pp. 4171–4186. Available: https://www.aclweb.org/anthology/N19-1423

38. Sun Y, Wang S, Li Y, Feng S, Chen X, Zhang H, et al. ERNIE: Enhanced Representation through Knowledge Integration. ArXiv. 2019;abs/1904.09223.

39. Anderson AJ, Kiela D, Binder JR, Fernandino L, Humphries CJ, Conant LL, et al. Deep Artificial Neural Networks Reveal a Distributed Cortical Network Encoding Propositional Sentence-Level Meaning. J Neurosci. 2021;41: 4100. doi:10.1523/JNEUROSCI.1152-20.2021

40. Hollenstein N, Renggli C, Glaus B, Barrett M, Troendle M, Langer N, et al. Decoding EEG Brain Activity for Multi-Modal Natural Language Processing. Front Hum Neurosci. 2021;15: 659410. doi:10.3389/fnhum.2021.659410

41. Toneva M, Wehbe L. Interpreting and improving natural-language processing (in machines) with natural language-processing (in the brain). In: Wallach H, Larochelle H, Beygelzimer A, Alché-Buc F d’, Fox E, Garnett R, editors. Advances in Neural Information Processing Systems. Curran Associates, Inc.; 2019. Available: https://proceedings.neurips.cc/paper/2019/file/749a8e6c231831ef7756db230b4359c8-Paper.pdf

42. Allen C, Hospedales T. Analogies Explained: Towards Understanding Word Embeddings. arXiv; 2019. Available: http://arxiv.org/abs/1901.09813

43. Levy O, Goldberg Y. Linguistic Regularities in Sparse and Explicit Word Representations. Proceedings of the Eighteenth Conference on Computational Natural Language Learning. Ann Arbor, Michigan: Association for Computational Linguistics; 2014. pp. 171–180. doi:10.3115/v1/W14-1618

44. Mikolov T, Sutskever I, Chen K, Corrado GS, Dean J. Distributed Representations of Words and Phrases and their Compositionality. NeurIPS Proc. 2013.

45. Pereira F, Lou B, Pritchett B, Ritter S, Gershman SJ, Kanwisher N, et al. Toward a universal decoder of linguistic meaning from brain activation. Nat Commun. 2018;9: 13. doi:10.1038/s41467-018-03068-4

46. Schaefer A, Kong R, Gordon EM, Laumann TO, Zuo XN, Holmes AJ, et al. Local- Global Parcellation of the Human Cerebral Cortex from Intrinsic Functional Connectivity MRI. Cereb Cortex. 2018;28: 3095–3114. doi:10.1093/cercor/bhx179

47. Satopaa V, Albrecht J, Irwin D, Raghavan B. Finding a “Kneedle” in a Haystack: Detecting Knee Points in System Behavior. 2011 31st International Conference on Distributed Computing Systems Workshops. Minneapolis, MN, USA: IEEE; 2011. pp. 166–171. doi:10.1109/ICDCSW.2011.20

48. Caucheteux C, Gramfort A, King J-R. Evidence of a predictive coding hierarchy in the human brain listening to speech. Nat Hum Behav. 2023;7: 430–441. doi:10.1038/s41562-022-01516-2

49. Huth AG, De Heer WA, Griffiths TL, Theunissen FE, Gallant JL. Natural speech reveals the semantic maps that tile human cerebral cortex. Nature. 2016;532: 453–458. doi:10.1038/nature17637

50. Millet J, Caucheteux C, Orhan P, Boubenec Y, Gramfort A, Dunbar E, et al. Toward a realistic model of speech processing in the brain with self-supervised learning. 36th Conf Neural Inf Process Syst NeurIPS 2022. 2022 [cited 16 Nov 2022]. Available: http://arxiv.org/abs/2206.01685

51. Esterman M, Tamber-Rosenau BJ, Chiu Y-C, Yantis S. Avoiding non- independence in fMRI data analysis: Leave one subject out. NeuroImage. 2010;50: 572–576. doi:10.1016/j.neuroimage.2009.10.092

52. Benjamini Y, Hochberg Y. Controlling the False Discovery Rate: A Practical and Powerful Approach to Multiple Testing. J R Stat Soc Ser B-Stat Methodol. 1995;57: 289–300. doi:10.1111/j.2517-6161.1995.tb02031.x

53. Nichols TE, Holmes AP. Nonparametric permutation tests for functional neuroimaging: A primer with examples. Hum Brain Mapp. 2002;15: 1–25. doi:10.1002/hbm.1058

54. Zhou C, Zwilling CE, Calhoun VD, Wang MY. Efficient Blockwise Permutation Tests Preserving Exchangeability. Int J Stat Med Res. 2014;3: 134–144. doi:10.6000/1929-6029.2014.03.02.8

55. Toneva M, Mitchell TM, Wehbe L. Combining computational controls with natural text reveals aspects of meaning composition. Nat Comput Sci. 2022;2: 745–757. doi:10.1038/s43588-022-00354-6

56. Alamia A, VanRullen R. Alpha oscillations and traveling waves: Signatures of predictive coding? PLOS Biol. 2019;17: 26. doi:10.1371/journal.pbio.3000487

57. Friston KJ. A theory of cortical responses. Philos Trans R Soc B-Biol Sci. 2005;360: 815–836. doi:10.1098/rstb.2005.1622

58. Chao ZC, Takaura K, Wang L, Fujii N, Dehaene S. Large-Scale Cortical Networks for Hierarchical Prediction and Prediction Error in the Primate Brain. Neuron. 2018;100: 1252-+. doi:10.1016/j.neuron.2018.10.004

59. Clark A. Whatever next? Predictive brains, situated agents, and the future of cognitive science. Behav Brain Sci. 05/10 ed. 2013;36: 181–204. doi:10.1017/S0140525X12000477

60. de Lange FP, Heilbron M, Kok P. How Do Expectations Shape Perception? Trends Cogn Sci. 2018;22: 764–779. doi:10.1016/j.tics.2018.06.002

61. Friston K. Does predictive coding have a future? Nat Neurosci. 2018;21: 1019– 1021. doi:10.1038/s41593-018-0200-7

62. Arnal LH, Giraud A-L. Cortical oscillations and sensory predictions. Trends Cogn Sci. 2012;16: 390–398. doi:10.1016/j.tics.2012.05.003

63. Bastos AM, Usrey WM, Adams RA, Mangun GR, Fries P, Friston KJ. Canonical Microcircuits for Predictive Coding. Neuron. 2012;76: 695–711. doi:10.1016/j.neuron.2012.10.038

64. Summerfield C, Egner T, Greene M, Koechlin E, Mangels J, Hirsch J. Predictive codes for forthcoming perception in the frontal cortex. Science. 2006;314: 1311– 1314. doi:10.1126/science.1132028

65. Finn ES, Corlett PR, Chen G, Bandettini PA, Constable RT. Trait paranoia shapes inter-subject synchrony in brain activity during an ambiguous social narrative. Nat Commun. 2018;9: 2043. doi:10.1038/s41467-018-04387-2

66. Whitney C, Huber W, Klann J, Weis S, Krach S, Kircher T. Neural correlates of narrative shifts during auditory story comprehension. Neuroimage. 2009;47: 360–366. doi:10.1016/j.neuroimage.2009.04.037

67. Zacks JM, Braver TS, Sheridan MA, Donaldson DI, Snyder AZ, Ollinger JM, et al. Human brain activity time-locked to perceptual event boundaries. Nat Neurosci. 2001;4: 651–655. doi:10.1038/88486

68. Friston KJ, Mechelli A, Turner R, Price CJ. Nonlinear Responses in fMRI: The Balloon Model, Volterra Kernels, and Other Hemodynamics. NeuroImage. 2000;12: 466–477. doi:10.1006/nimg.2000.0630

69. Zeidman P, Jafarian A, Corbin N, Seghier ML, Razi A, Price CJ, et al. A guide to group effective connectivity analysis, part 1: First level analysis with DCM for fMRI. Neuroimage. 2019;200: 174–190. doi:10.1016/j.neuroimage.2019.06.031

70. Franklin NT, Norman KA, Ranganath C, Zacks JM, Gershman SJ. Structured Event Memory: A neuro-symbolic model of event cognition. Psychol Rev. 2020/04/01 ed. 2020;127: 327–361. doi:10.1037/rev0000177

71. Armeni K, Willems RM, van den Bosch A, Schoffelen JM. Frequency-specific brain dynamics related to prediction during language comprehension. Neuroimage. 2019;198: 283–295. doi:10.1016/j.neuroimage.2019.04.083

72. Grisoni L, Miller TM, Pulvermüller F. Neural Correlates of Semantic Prediction and Resolution in Sentence Processing. J Neurosci. 2017;37: 4848–4858. doi:10.1523/jneurosci.2800-16.2017

73. Grisoni L, Mohr B, Pulvermüller F. Prediction mechanisms in motor and auditory areas and their role in sound perception and language understanding. NeuroImage. 2019;199: 206–216. doi:10.1016/j.neuroimage.2019.05.071

74. Lopopolo A, Frank SL, van den Bosch A, Willems RM. Using stochastic language models (SLM) to map lexical, syntactic, and phonological information processing in the brain. PLOS One. 2017;12. doi:10.1371/journal.pone.0177794

75. Kutas M, Federmeier KD. Thirty Years and Counting: Finding Meaning in the N400 Component of the Event-Related Brain Potential (ERP). In: Fiske ST, Schacter DL, Taylor SE, editors. Annual Review of Psychology. 2011. pp. 621– 647. doi:10.1146/annurev.psych.093008.131123

76. Armeni K, Willems RM, Frank SL. Probabilistic language models in cognitive neuroscience: Promises and pitfalls. Neurosci Biobehav Rev. 2017;83: 579–588. doi:10.1016/j.neubiorev.2017.09.001

77. Miller EK. The prefrontal cortex and cognitive control. Nat Rev Neurosci. 2000;1: 59–65. doi:10.1038/35036228

78. Wood JN, Grafman J. Human prefrontal cortex: Processing and representational perspectives. Nat Rev Neurosci. 2003;4: 139–147. doi:10.1038/nrn1033

79. Chen J, Leong YC, Honey CJ, Yong CH, Norman KA, Hasson U. Shared memories reveal shared structure in neural activity across individuals. Nat Neurosci. 2017;20: 115–125. doi:10.1038/nn.4450

80. Yeshurun Y, Swanson S, Simony E, Chen J, Lazaridi C, Honey CJ, et al. Same Story, Different Story: The Neural Representation of Interpretive Frameworks. Psychol Sci. 2017;28: 307–319. doi:10.1177/0956797616682029

81. Hasson U, Chen J, Honey CJ. Hierarchical process memory: memory as an integral component of information processing. Trends Cogn Sci. 2015;19: 304–313. doi:10.1016/j.tics.2015.04.006

82. Schacter DL, Addis DR, Buckner RL. Episodic Simulation of Future Events: Concepts, Data, and Applications. Ann N Y Acad Sci. 2008;1124: 39–60. doi:10.1196/annals.1440.001

83. Preminger S, Harmelech T, Malach R. Stimulus-free thoughts induce differential activation in the human default network. NeuroImage. 2011;54: 1692–1702. doi:10.1016/j.neuroimage.2010.08.036

84. Gilbert DT, Wilson TD. Prospection: Experiencing the Future. Science. 2007;317: 1351–1354. doi:10.1126/science.1144161

85. Ferstl EC, Neumann J, Bogler C, von Cramon DY. The extended language network: A meta-analysis of neuroimaging studies on text comprehension. Hum Brain Mapp. 2008;29: 581–593. doi:10.1002/hbm.20422

86. Hamilton LS, Huth AG. The revolution will not be controlled: natural stimuli in speech neuroscience. Lang Cogn Neurosci. 2020;35: 573–582. doi:10.1080/23273798.2018.1499946

87. Hasson U, Egidi G, Marelli M, Willems RM. Grounding the neurobiology of language in first principles: The necessity of non-language-centric explanations for language comprehension. Cognition. 2018;180: 135–157. doi:10.1016/j.cognition.2018.06.018

88. Sridharan D, Levitin DJ, Chafe CH, Berger J, Menon V. Neural Dynamics of Event Segmentation in Music: Converging Evidence for Dissociable Ventral and Dorsal Networks. Neuron. 2007;55: 521–532. doi:10.1016/j.neuron.2007.07.003

89. Bullmore E, Sporns O. The economy of brain network organization. Nat Rev Neurosci. 2012;13: 336–349. doi:10.1038/nrn3214

90. Sporns O. The Non-Random Brain: Efficiency, Economy, and Complex Dynamics. Front Comput Neurosci. 2011;5. doi:10.3389/fncom.2011.00005

91. Kamide Y, Altmann GTM, Haywood SL. The time-course of prediction in incremental sentence processing: Evidence from anticipatory eye movements. J Mem Lang. 2003;49: 133–156. doi:10.1016/S0749-596X(03)00023-8

92. Sedivy JC, K. Tanenhaus M, Chambers CG, Carlson GN. Achieving incremental semantic interpretation through contextual representation. Cognition. 1999;71: 109–147. doi:10.1016/S0010-0277(99)00025-6

93. Payne BR, Lee C, Federmeier KD. Revisiting the incremental effects of context on word processing: Evidence from single-word event-related brain potentials. Psychophysiology. 2015;52: 1456–1469. doi:10.1111/psyp.12515

94. Leon-Cabrera P, Flores A, Rodriguez-Fornells A, Moris J. Ahead of time: Early sentence slow cortical modulations associated to semantic prediction. Neuroimage. 2019;189: 192–201. doi:10.1016/j.neuroimage.2019.01.005

95. Oldfield RC. The assessment and analysis of handedness: the Edinburgh inventory. Neuropsychologia. 1971;9: 97–113. doi:10.1016/0028-3932(71)90067-4

96. Liu LF, Zhang YX, Zhou Q, Garrett DD, Lu CM, Chen AT, et al. Auditory- Articulatory Neural Alignment between Listener and Speaker during Verbal Communication. Cereb Cortex. 2020;30: 942–951. doi:10.1093/cercor/bhz138

97. Silbert LJ, Honey CJ, Simony E, Poeppel D, Hasson U. Coupled neural systems underlie the production and comprehension of naturalistic narrative speech. Proc Natl Acad Sci U S A. 2014;111: E4687–E4696. doi:10.1073/pnas.1323812111

98. Stephens GJ, Silbert LJ, Hasson U. Speaker-listener neural coupling underlies successful communication. Proc Natl Acad Sci U S A. 2010;107: 14425–14430. doi:10.1073/pnas.1008662107

99. Nicholas LE, Brookshire RH. Consistency of the Effects of Rate of Speech on Brain-Damaged Adults’ Comprehension of Narrative Discourse. J Speech Hear Res. 1986;29: 462–470. doi:10.1044/jshr.2904.462

100. Yan CG, Wang XD, Zuo XN, Zang YF. DPABI: Data Processing & Analysis for (Resting-State) Brain Imaging. Neuroinformatics. 2016;14: 339–351. doi:10.1007/s12021-016-9299-4

101. Friston KJ, Williams S, Howard R, Frackowiak RSJ, Turner R. Movement- Related effects in fMRI time-series: Movement Artifacts in fMRI. Magn Reson Med. 1996;35: 346–355. doi:10.1002/mrm.1910350312

102. Qiao Y, Xiong C, Liu Z, Liu Z. Understanding the Behaviors of BERT in Ranking. arXiv; 2019. Available: http://arxiv.org/abs/1904.07531

103. Alzahrani E, Jololian L. How Different Text-Preprocessing Techniques using the Bert Model Affect the Gender Profiling of Authors. Advances in Machine Learning. Academy and Industry Research Collaboration Center (AIRCC); 2021. pp. 01–08. doi:10.5121/csit.2021.111501

104. Vaswani A, Shazeer N, Parmar N, Uszkoreit J, Jones L, Gomez AN, et al. Attention Is All You Need. In: Guyon I, Luxburg UV, Bengio S, Wallach H, Fergus R, Vishwanathan S, et al., editors. Advances in Neural Information Processing Systems 30. La Jolla: Neural Information Processing Systems (Nips); 2017. Available: ://WOS:000452649406008

105. Huang J, Tang D, Zhong W, Lu S, Shou L, Gong M, et al. WhiteningBERT: An Easy Unsupervised Sentence Embedding Approach. arXiv; 2021. Available: http://arxiv.org/abs/2104.01767

106. Yu L, Ettinger A. Assessing Phrasal Representation and Composition in Transformers. arXiv; 2020. Available: http://arxiv.org/abs/2010.03763

107. Pedregosa F, Varoquaux G, Gramfort A, Michel V, Thirion B, Grisel O, et al. Scikit-learn: Machine Learning in Python. J Mach Learn Res. 2011;12: 2825– 2830.

108. Jha R, Mihata K. On Geodesic Distances and Contextual Embedding Compression for Text Classification. Mexico City, Mexico: Association for Computational Linguistics; 2021. pp. 144–149. doi:10.18653/v1/2021.textgraphs-1.15

109. Tenenbaum JB, de Silva V, Langford JC. A global geometric framework for nonlinear dimensionality reduction. Science. 2000/12/23 ed. 2000;290: 2319–23. doi:10.1126/science.290.5500.2319

110. Cai Q, Brysbaert M. SUBTLEX-CH: Chinese Word and Character Frequencies Based on Film Subtitles. PLOS One. 2010;5: 8. doi:10.1371/journal.pone.0010729

111. New B, Brysbaert M, Veronis J, Pallier C. The use of film subtitles to estimate word frequencies. Appl Psycholinguist. 2007;28: 661–677. doi:10.1017/s014271640707035x

112. Abraham A, Pedregosa F, Eickenberg M, Gervais P, Mueller A, Kossaifi J, et al. Machine learning for neuroimaging with scikit-learn. Front Neuroinformatics. 2014;8. doi:10.3389/fninf.2014.00014

113. Xia M, Wang J, He Y. BrainNet Viewer: A Network Visualization Tool for Human Brain Connectomics. Csermely P, editor. PLoS ONE. 2013;8: e68910. doi:10.1371/journal.pone.0068910

114. Friston KJ. Waves of prediction. PLOS Biol. 2019;17: e3000426. doi:10.1371/journal.pbio.3000426

115. Summerfield C, Egner T, Mangels J, Hirsch J. Mistaking a house for a face: Neural correlates of misperception in healthy humans. Cereb Cortex. 2006;16: 500–508. doi:10.1093/cercor/bhi129

116. Summerfield C, Trittschuh EH, Monti JM, Mesulam MM, Egner T. Neural repetition suppression reflects fulfilled perceptual expectations. Nat Neurosci. 2008;11: 1004–1006. doi:10.1038/nn.2163

117. Brodbeck C, Hong LE, Simon JZ. Rapid Transformation from Auditory to Linguistic Representations of Continuous Speech. Curr Biol. 2018;28: 3976-+. doi:10.1016/j.cub.2018.10.042

118. Nguyen HV, Bai L. Cosine Similarity Metric Learning for Face Verification. In: Kimmel R, Klette R, Sugimoto A, editors. Computer Vision – ACCV 2010. Berlin, Heidelberg: Springer Berlin Heidelberg; 2011. pp. 709–720. doi:10.1007/978-3-642-19309-5_55

119. Zhang L, Wang L, Yang J, Qian P, Wang X, Qiu X, et al. Can computers understand words like humans do? Comparable semantic representation in neural and computer systems. bioRxiv. 2020; 843896. doi:10.1101/843896

120. Stephan KE, Weiskopf N, Drysdale PM, Robinson PA, Friston KJ. Comparing hemodynamic models with DCM. NeuroImage. 2007;38: 387–401. doi:10.1016/j.neuroimage.2007.07.040

121. Seabold S, Perktold J. Statsmodels: Econometric and Statistical Modeling with Python. Austin, Texas; 2010. pp. 92–96. doi:10.25080/Majora-92bf1922-011

